# Spatial and temporal coordination of signaling pathways in tissue differentiation: developmental atlas of protein expression during zebra finch beak maturation

**DOI:** 10.64898/2025.12.17.695020

**Authors:** Renée A. Duckworth, Sarah E. Britton, Cody A. Lee, Kathryn C. Chenard, Alexander V. Badyaev

**Affiliations:** Department of Ecology and Evolutionary Biology, University of Arizona, Tucson, AZ USA; Department of Ecology, Behavior and Evolution and; School of Medicine, University of California San Diego, La Jolla, CA, USA; Department of Integrative Biology, University of Texas at Austin, Austin, TX USA

**Keywords:** craniofacial development, chondrogenesis, osteogenesis, immunohistochemistry, mesenchymal stem cells

## Abstract

**Background:** Morphogenesis depends on spatial and temporal coordination of signaling pathways, yet the colocalization of proteins across pathways remains poorly understood. Here we examine cellular and histological localization of regulatory proteins forming core craniofacial developmental pathways during beak morphogenesis of the zebra finch (*Taeniopygia guttata*).

**Results:** We present an atlas of spatiotemporal coexpression of β-catenin, Bmp4, CaM, Dkk3, Fgf8, Ihh, Tgfβ2, and Wnt4 across embryonic stages HH29-42 revealing both established and novel patterns of expression. Overall, in the earliest stages (HH29-32), most proteins show broad and overlapping expression across epithelial and mesenchymal tissues. By stage HH36, expression becomes increasingly compartmentalized, with pronounced differentiation among tissue types. Notably, at later stages, proteins showed tissue-specific distributions in boundary versus core regions of chondrogenic and osteogenic domains indicating coordinated cross-pathway patterning during cartilage and bone formation.

**Conclusions:** Osteogenesis in the zebra finch beak is organized by coordinated signaling between boundary-associated cells and differentiating cores, with cross-pathway feedback establishing bone and cartilage differentiation while maintaining boundaries. Our results corroborated core elements of craniofacial signaling dynamics, while revealing unexpected subcellular localization for several proteins that showed regulatory complexity not captured by prior transcript-level maps. This atlas provides a protein-level baseline for comparative and mechanistic studies of beak morphogenesis.

## Introduction

During craniofacial development, signaling proteins exhibit complex and dynamic expression patterns that integrate positional information and coordinate tissue differentiation^1–4^. The functions of these signals ultimately require their physical assembly, necessitating subcellular and within-tissue colocalization. In particular, distinguishing membranal, cytoplasmic, and nuclear compartmentalization of protein complexes is crucial for understanding pathway activation states and the directionality of communication at the tissue level^5^. For example, nuclear localization of transcriptional regulators could indicate active signaling, whereas cytoplasmic localization suggests ligand production or initiation of intracellular signaling cascades^6–8^. By placing subcellular localizations in a spatial and temporal context, we can gain insight into the flow and distribution of information among cells as well as the functional importance of expression overlap for tissue differentiation. Further, mapping the colocalization and compartmentalization of regulatory proteins across multiple signaling pathways allows for insight into the temporal and spatial activation of protein networks and their interactions in morphogenesis and evolution.

The development of the avian beak is a model for studies of evolutionary divergence in vertebrates and exploration of craniofacial abnormalities in humans^9,10^. Both the evolutionary diversification of beak morphology and its relevance to human craniofacial disorders stem from a shared developmental architecture governed by four evolutionarily conserved signaling pathways: wingless type protein (Wnt), bone morphogenetic protein (Bmp; part of the transcription growth factor, Tgfβ, superfamily), fibroblast growth factor (Fgf), and hedgehog (Hh) signaling^11–15^. Here, we examine the cellular localization, developmental timing, and tissue-specific expression of eight key regulatory proteins from these pathways that have been well characterized in early stages of avian beak development^16–22^: β-catenin, bone morphogenic protein 4 (Bmp4), calmodulin (CaM), dickkopf homolog 3 (Dkk3), fibroblast growth factor 8 (Fgf8), Indian hedgehog (Ihh), transforming growth factor beta 2 (Tgfβ2), and wingless type 4 (Wnt4).

Wnt signaling regulates cell polarity, proliferation, and fate specification and interacts extensively with the other pathways integrating signals that orchestrate tissue patterning^23^ (Figure 1). β-catenin and Wnt4 capture canonical and non-canonical components of the Wnt pathway ^19,24^, respectively, while Dkk3 serves as a context-dependent Wnt modulator^25,26^ (Figure 1). Bmp4 and Tgfβ2, both members of the TGF-β superfamily, represent key regulators of mesenchymal differentiation and epithelial–mesenchymal signaling^27–29^. FGF signaling, via four receptor subtypes, fine-tunes growth and differentiation of skeletal progenitors through feedback loops that ensure spatial and temporal precision^30–32^. Fgf8 is a central mediator of epithelial patterning and mesenchymal proliferation within the frontonasal region^33,34^. Hh signaling, primarily through Sonic and Indian Hedgehog (Shh and Ihh, respectively), maintains neural crest survival and coordinates epithelial–mesenchymal interactions essential for jaw and midline formation^35–37^. Finally, CaM, a calcium-binding transducer interacts with multiple of these pathways (Figure 1) and has an integrative role in osteoblast growth and differentiation^38^ and beak shape evolution^39^.

**Figure 1.**
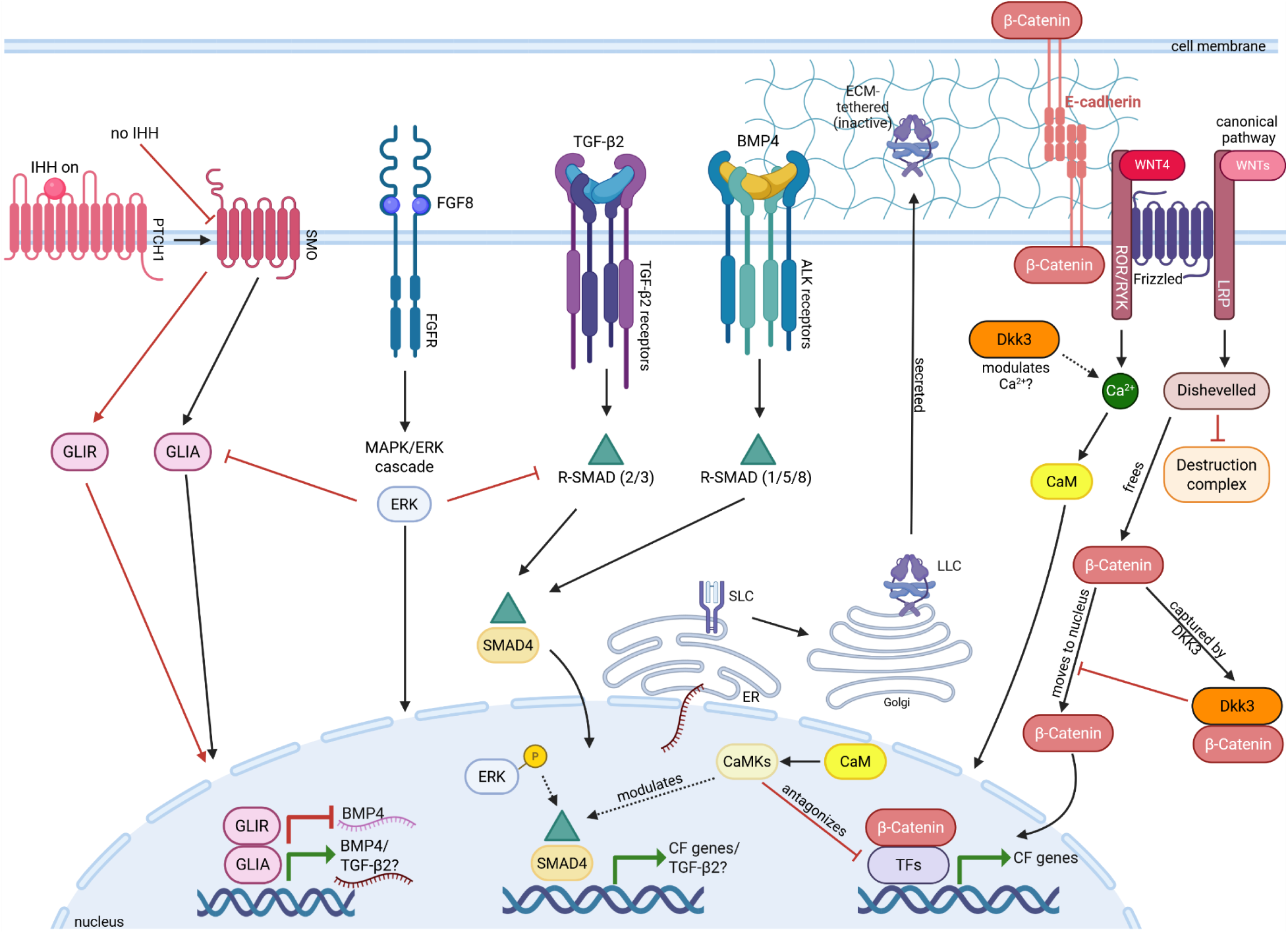
Summary of known and potential interactions of eight focal proteins in craniofacial development. HH signaling (far left), is initiated when IHH binds to cell surface receptor Patched 1 (PTCH1), activating Smoothened (SMO) which, in turn, initiates a signaling cascade activating Gli activator proteins (GLIA) which are key transcription factors for craniofacial (CF) genes, including BMP4 and TGFβ2. In the absence of IHH ligand, SMO is blocked and the Gli repressor (GLIR) is produced blocking gene expression. BMP4 and TGFβ2 each initiate signaling cascades involving different sets of R-SMAD proteins that form a complex with SMAD4. This complex translocates to the nucleus where it recruits other co-activators or repressors (not shown) and binds DNA to activate (and sometimes repress) target CF genes, including in some cases TGFβ2. FGF8 signaling can block IHH, TGFβ2, and BMP4 pathways as it activates the MAPK/ERK pathway and ERK can inhibit both GLIA and R-SMADs from reaching the nucleus. However, in some developmental contexts, ERK can enhance gene transcription or shift it to new target genes by phosphorylating R-SMADs in the nucleus. Unlike the other secreted ligands, TGFβ2 is translated as a pro-peptide that dimerizes in the endoplasmic reticulum (ER) and forms the small latent complex (SLC), which then binds latent TGFβ-binding protein in the Golgi to form the large latent complex (LLC). The LLC is secreted and tethered to the ECM in an inactive state until mechanically or proteolytically activated. WNT4 (far right) mainly acts as a non-canonical Wnt ligand by binding to a Frizzled/ROR- or Frizzled/RYK-type receptor complex to activate Ca²⁺ signaling which elevates intracellular Ca²⁺ and activates calmodulin (CaM) and downstream kinases (CamKs) either in the cytoplasm (not shown) or the nucleus. These kinases, in turn, provide context-dependent modulation of Smads and other CF-relevant transcription factors (TF). In parallel, a separate canonical Wnt pathway activates the LRP-Frizzled-Dishevelled axis, blocking the destruction complex and stabilizing cytoplasmic β-catenin. This enables its nuclear translocation to regulate craniofacial (CF) genes with TCF/LEF (T cell factor/lymphoid enhancer factor family; not shown) and other transcription factors. Non-canonical WNT4/Ca²⁺/CaM signaling can antagonize canonical output by inhibiting TCF/β-catenin-dependent transcriptional activity. Dkk3 is a context-dependent modulator of this axis, as it has been proposed to modulate Ca²⁺ production although the mechanism is still under investigation. DKK3 can also capture cytoplasmic β-catenin, limiting its nuclear entry and further impacting canonical Wnt activity. In the absence of canonical Wnt signaling, a pool of β-catenin is bound to E-cadherin at adherens junctions in the cell membrane, contributing to cell–cell adhesion and sequestering β-catenin away from the nucleus.

Together these factors and their pathways form an interconnected regulatory network that orchestrates craniofacial patterning during development (Figure 1). For example, Fgf8 activates the MAPK–ERK (mitogen-activated protein kinases-extracellular signal-regulated kinases) pathway, and sustained ERK activity suppresses both the Bmp4 Smad cascade as well as Ihh–Gli signaling^40–42^. Bmp4 and Tgfβ2 activate distinct Smad pathways that converge on shared gene regulatory networks, producing synergistic or antagonistic effects on target genes depending on whether they compete for Smad4 or their Smad complexes co-occupy enhancer or promoter regions^43,44^. Wnt4 signaling interacts with all of these pathways by impacting β-catenin dynamics, often inhibiting the canonical Wnt pathway and redirecting β-catenin to the cell membrane, effectively preventing it from reaching the nucleus (Figure 1). However, in some tissue contexts^26^, it can also regulate the canonical Wnt pathway and free β-catenin from its destruction complex promoting its movement to the nucleus, where β-catenin co-binds with various transcription factors, including Bmp- and Tgfβ-activated Smads, to regulate numerous craniofacial genes^7,19,23,26^. In turn, Dkk3 can shift Wnt responses toward non-canonical, Ca²⁺-dependent pathways, engaging CaM signaling^45,46^ and it can also prevent β-catenin from entering the nucleus^47^. The balance among these pathways is highly context-dependent: the same signaling input can produce opposite transcriptional outcomes depending on Smad availability, Gli activator–repressor ratios, or competition between canonical and non-canonical Wnt branches^48–50^. Yet, how this balance shifts within the context of avian beak morphogenesis is not well characterized, and mapping protein colocalization across stages is needed to understand how pathway interactions might influene diversity in craniofacial development.

In this study, we generate an atlas of zebra finch (*Taeniopygia guttata*) beak ontogeny across five developmental stages (Hamburger-Hamilton, HH, 29/30, 32, 36, 39, and 42) that span key late-development transitions in tissue differentiation and bone and cartilage formation. The zebra finch is an emerging model system in developmental biology and neurobiology^51–54^, with high quality genomic resources^55^ and well-described developmental staging^56^. Moreover, zebra finches are particularly well suited as a model for developmental studies of the avian beak because their conical beaks share key functional requirenments with other well-studied finch and seed-eating birds^57,58^. Here, we use immunohistochemistry with chromogenic labeling and nuclear counterstaining, to map both the spatiotemporal expression and subcellular localization of focal proteins across tissues.

## Results

### Temporal dynamics of protein expression and subcellular localization across tissues

Over developmental progression, diffuse, mesenchymal expression of proteins transitioned to localized, pathway-specific patterns of expression, corresponding to morphogenetic patterning and structural maturation of the beak (Figures 2-12). In the earliest stages (HH29-32), most proteins showed broad and overlapping expression across epithelial and mesenchymal tissues (Figures 3-6). By stage HH36, expression became increasingly compartmentalized, with pronounced differentiation among tissue types (Figures 7, 8). Strong expression of several proteins in myogenic, chondrogenic, and nasal epithelial regions at this stage indicated functionally distinct and tissue-specific signaling environments. In later stages (HH39–42; Figures 9-12), expression of most proteins was spatially restricted to osteogenic, perichondrogenic, and myogenic domains.

**Figure 2.**
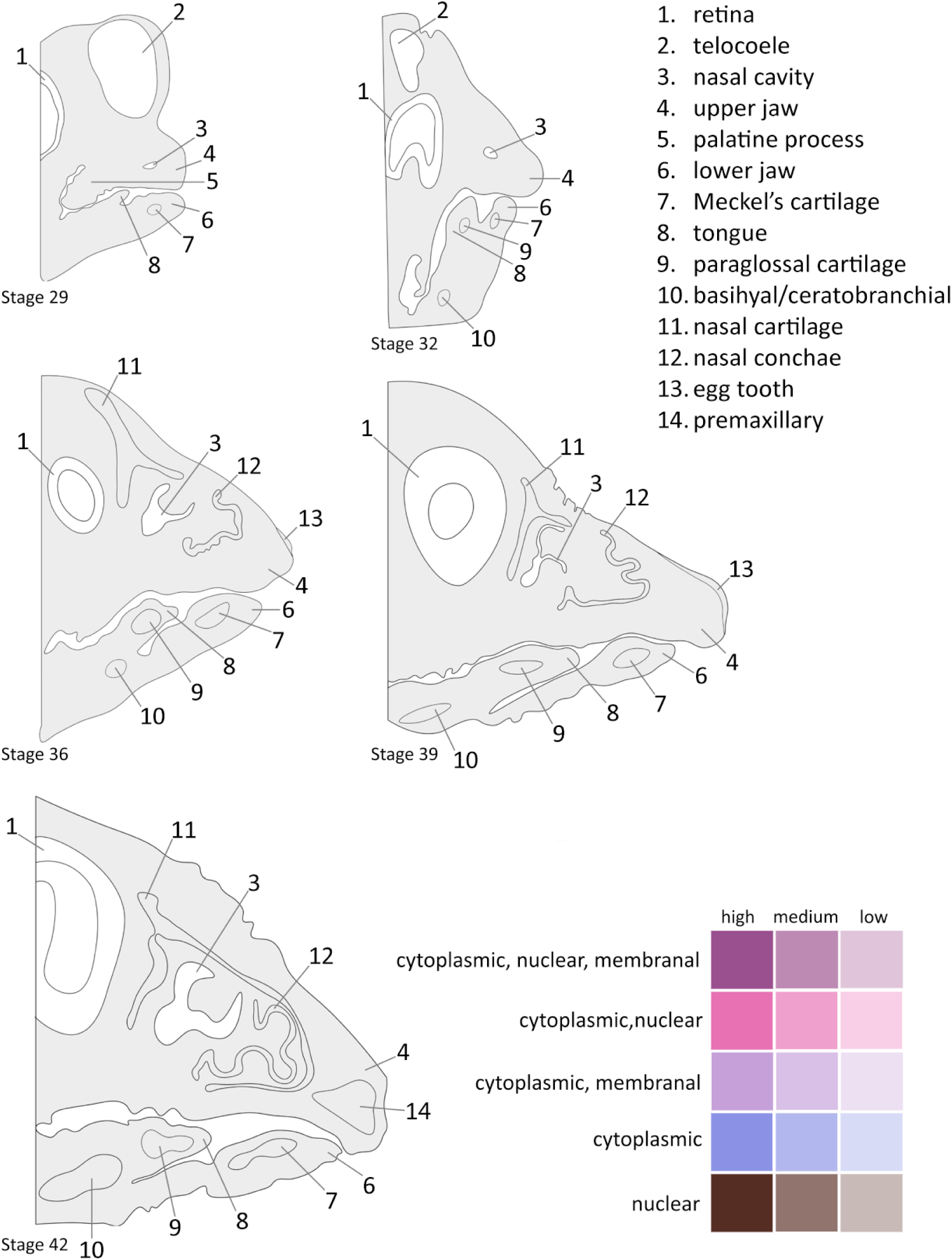
Consensus illustrations of mid-sagittal sections of 5 developmental stages traced from prototypical embryos used in this study with major anatomical features labeled. Key shows color scheme used to denote cellular localization of expression as well as expression intensity in Figures 3-12. Nuclear expression includes perinuclear.

**Figure 3.**
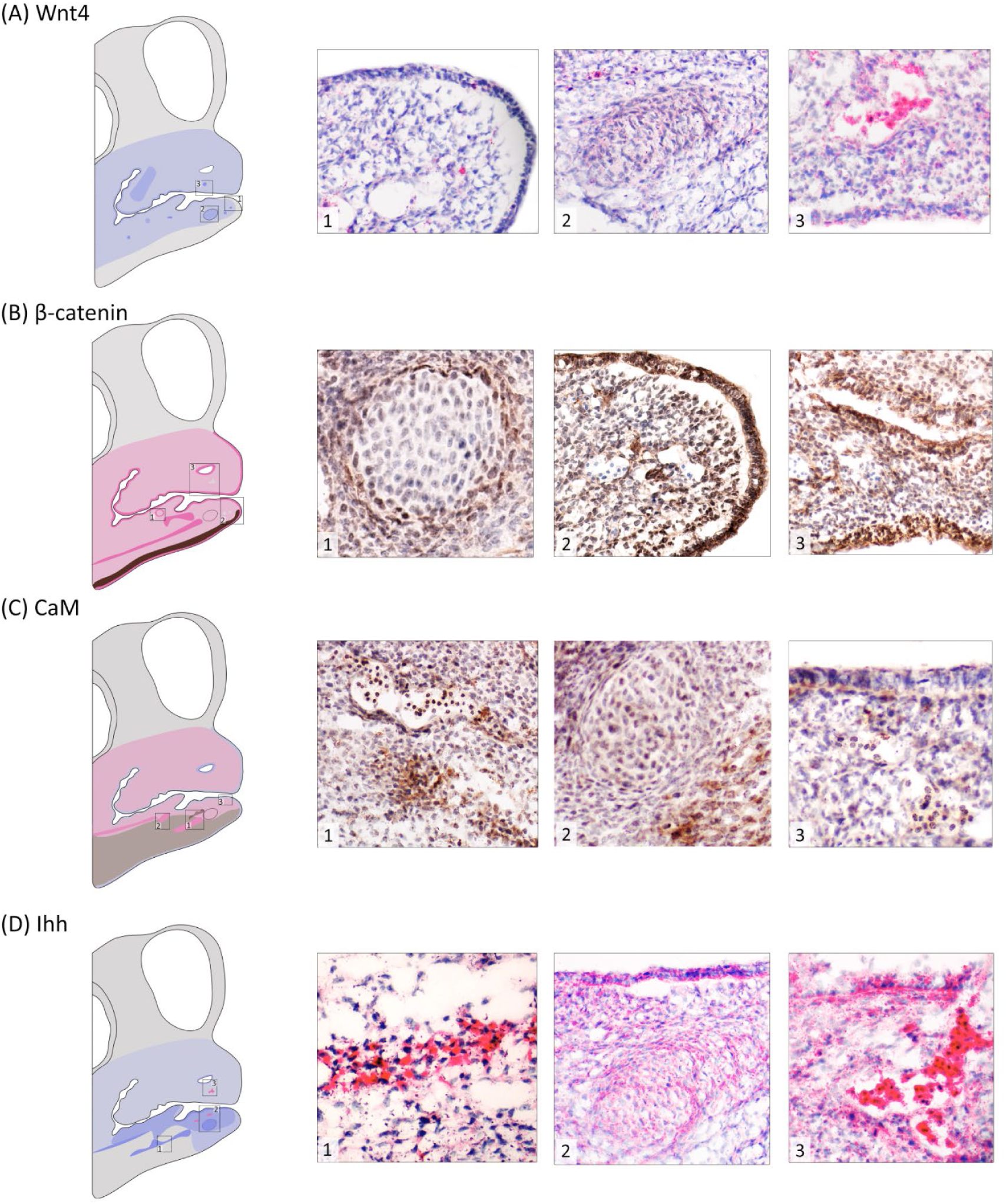
Wnt4, β-catenin, CaM, and Ihh expression at stage HH29/30. (A) **Wnt4** (red stain) shows relatively weak cytoplasmic expression overall with (1) no expression at the tip and ventral portion of the mandibular prominence and (2) moderate expression in the Meckel’s condensation. (3) Strong cytoplasmic and perinuclear expression occurred in hematopoietic cells. (B) **β-catenin** expression (2^nd^ panel, brown stain) is strong in both jaws with (1) the weakest expression in pre-cartilage condensations (paraglossal shown) which are surrounded by cells showing strong nuclear expression. (2) Nuclear and cytoplasmic expression in the surface epithelium of both jaws (lower mandibular tip shown) with expression limited to nuclear in the ventral portion of the mandibular prominence. There is no expression in hematopoietic cells. (3) Strong nuclear and cytoplasmic expression in the nasal and oral epithelium of the maxillary prominence. (C) **CaM** (brown stain) shows moderate nuclear and cytoplasmic expression in both jaws with (1, 2) stronger expression in potentially myogenic areas adjacent to pre-cartilage condensations. There is strong nuclear expression in hematopoietic tissue. (3) Epithelial expression is cytoplasmic with scattered nuclear expression in the underlying mesenchyme. (D) **Ihh** (red stain) shows overall weak expression in the maxillary prominence, and is stronger, albeit more localized to specific regions, in the mandibular prominence. (1) There is a stripe of strong cytoplasmic expression in likely pre-myogenic tissue along the center of the mandibular prominence and (2) moderate expression in the oral epithelium that matches underlying expression in the Meckel’s condensation. (3) Expression in the maxillary prominence is overall weak with the exception of strong cytoplasmic expression in hematopoietic tissue and at the edge of the nasal cavity. See Figure 2 for key to anatomical features and expression patterns. Photos here and throughout 40x.

Two exceptions to this generalized progression were β-catenin and Tgfβ2 which showed strong and widespread expression across all stages. β-catenin showed strong nuclear expression in early stages before shifting to membranous localization in the epithelium at later stages (e.g. Figure 9B-1), while Tgfβ2 was expressed perinuclearly within chondrogenic condensations (e.g. Figure 6C-1,4). Expression of Fgf8 was spatially restricted at HH29/30 (Figure 4) but underwent a major expansion at stage HH32 (Figure 6), particularly in the upper beak, followed by gradual restriction to epithelial and skeletal boundaries by stage HH42 (Figure 12). Ihh, Fgf8 and Tgfβ2 localized strongly to osteogenic fronts, cartilage, and muscle at mid to late stages, with Ihh, particularly, delineating areas of intramembranous ossification (Figures 11, 12). CaM, in contrast, showed relatively weak and scattered expression by stage HH42, with strongest expression in muscle and epithelial cells (Figure 11). Dkk3 showed strong expression at stage HH29/30 in areas surrounding condensations that would later form cartilage (Figure 4) while Wnt4 expression was stronger within condensations and absent without (e.g. Figures 3, 5). Thus, early broad expression of regulatory proteins became increasingly compartmentalized, aligning with tissue-specific differentiation and maturation of skeletal, muscular, and epithelial domains in the beak.

**Figure 4.**
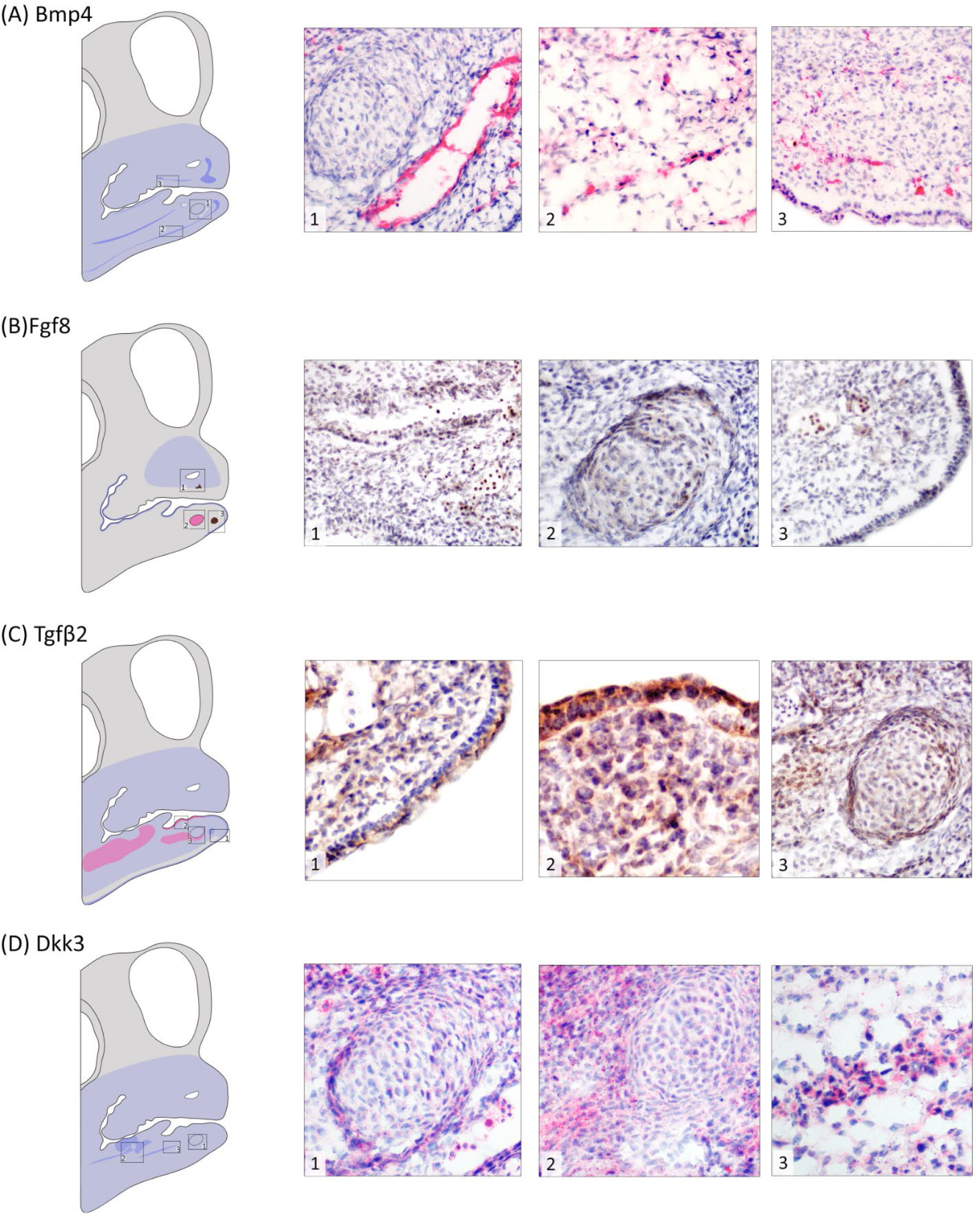
Bmp4, Fgf8, TGF-β, and Dkk3 expression at stage HH29/30. (A) **Bmp4** (red stain) shows relatively weak cytoplasmic expression overall with (1) with strong expression concentrated around blood vessels and in stripes of mesenchymal tissue in both the (2) lower and (3) upper ventral beak margins. Spatial extent of (B) **Fgf8** expression (brown stain) is limited in both jaws. (1) Maxillary prominence shows weak cytoplasmic expression around the nasal cavity with strong nuclear expression only in hematopoietic cells. (2) Meckel’s cartilage shows weak cytoplasmic expression with areas of strong perinuclear expression, especially at the periphery. (3) Weak cytoplasmic expression in the mesenchyme and surface epithelium. (C) **TGF-β** (brown stain) shows relatively weak cytoplasmic expression in the upper and lower beak prominences. (1) Weak cytoplasmic expression in mesenchyme shifts to moderately strong around developing vascular opening, no expression in hematopoietic cells, and moderately strong staining in surface epithelium of the mandibular prominence. (2) Strong cytoplasmic and perinuclear staining in the oral epithelium of the lower beak prominence. Meckel’s condensation shows weak cytoplasmic staining with moderate perinuclear expression in the perichondrogenic area and in an adjacent premyogenic area. (D) **Dkk3** (red stain) shows overall weak cytoplasmic expression in both prominences with regions of stronger expression in the mandibular prominence. (1) Elevated perichondrogenic expression for the Meckel’s condensation and moderate expression in hematopoietic cells. (2) Moderate expression in the mesenchyme surrounding the basihyal condensation with only weak expression in the condensation itself. (3) Moderate expression in a stripe of mesenchymal tissue in the ventral margin of the mandibular process. See Figure 2 for key to anatomical features and expression patterns.

**Figure 5.**
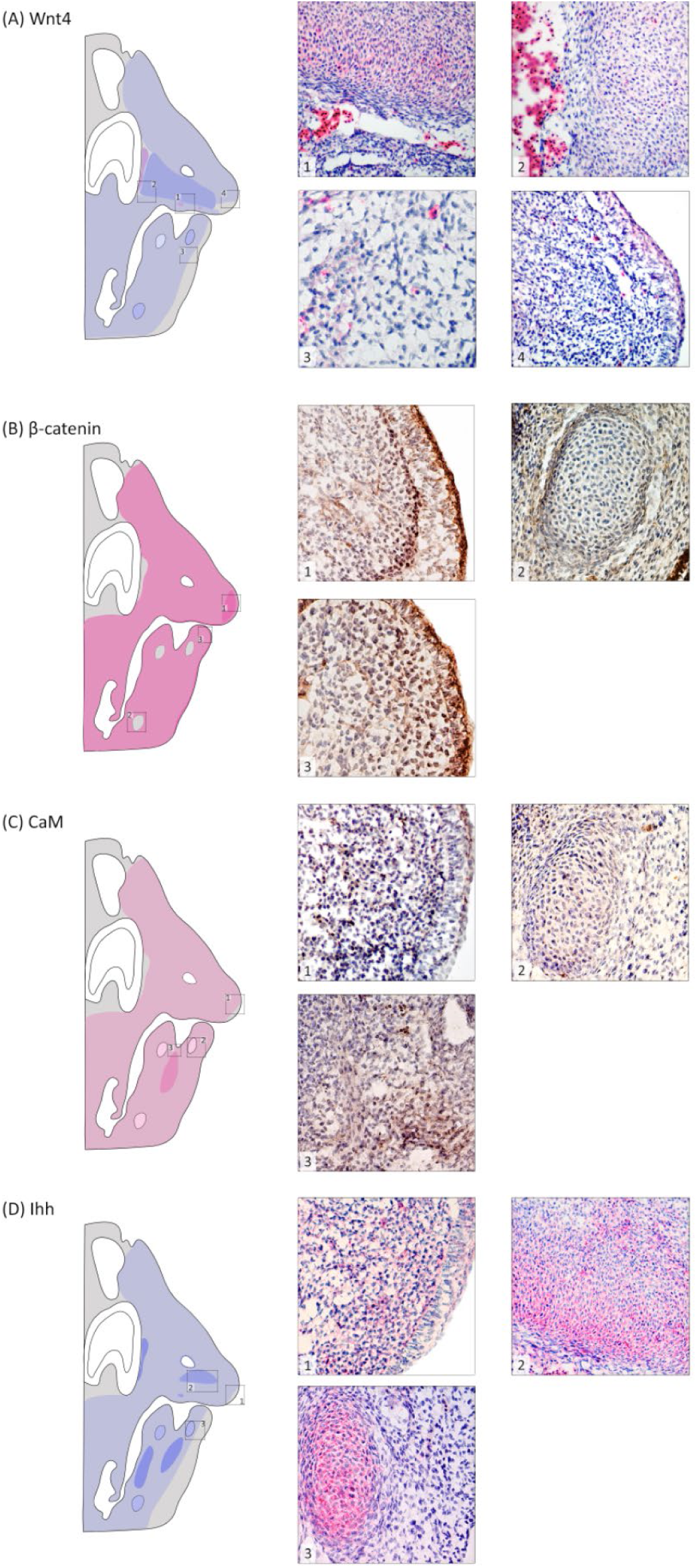
Wnt4, β-catenin, CaM, and Ihh expression at stage HH32. (A) **Wnt4** shows relatively weak cytoplasmic expression overall with (1,2) moderate expression in a large condensation of the maxillary prominence and strong expression in hematopoietic cells. (3) Weak expression in mandibular prominence with no expression along its ventral edge, including in the surface epithelium. (4) No expression in mesenchyme of maxillary prominence at the tip, however, surface epithelium shows weak expression. (B) **β-catenin** shows strong cytoplasmic and nuclear expression in both jaws with (1) the strongest nuclear expression just under the surface epithelium of the beak tip. (2) Expression is largely absent in the basihyal cartilage of the mandibular prominence but is present in some nuclei of perichondrogenic cells. (3) Strong nuclear and cytoplasmic expression in the surface epithelium of the mandibular prominence, with particularly strong nuclear expression in adjacent mesenchymal cells and elevated cytoplasmic expression surrounding developing vasculature. No expression in hematopoietic cells. (C) **CaM** shows weak nuclear and cytoplasmic expression in both jaws with (1) absence of expression at the beak tip. (2) Weak, mostly cytoplasmic, expression in the emerging Meckel’s cartilage with some scattered perinuclear expression. (3) Moderate expression in a region of mesenchyme just below the tongue which is likely premyogenic. (D) **Ihh** shows overall weak cytoplasmic expression in both jaws. (1) Expression is largely absent in the epithelium at the tip of the maxillary prominence. (2) Moderate to strong expression in the central condensation of the maxillary prominence. (3) Strong expression in the Meckel’s cartilage with notable absence of expression on the ventral margin of the mandibular prominence. See Figure 2 for key to anatomical features and expression patterns. Stain colors as in Figure 3.

**Figure 6.**
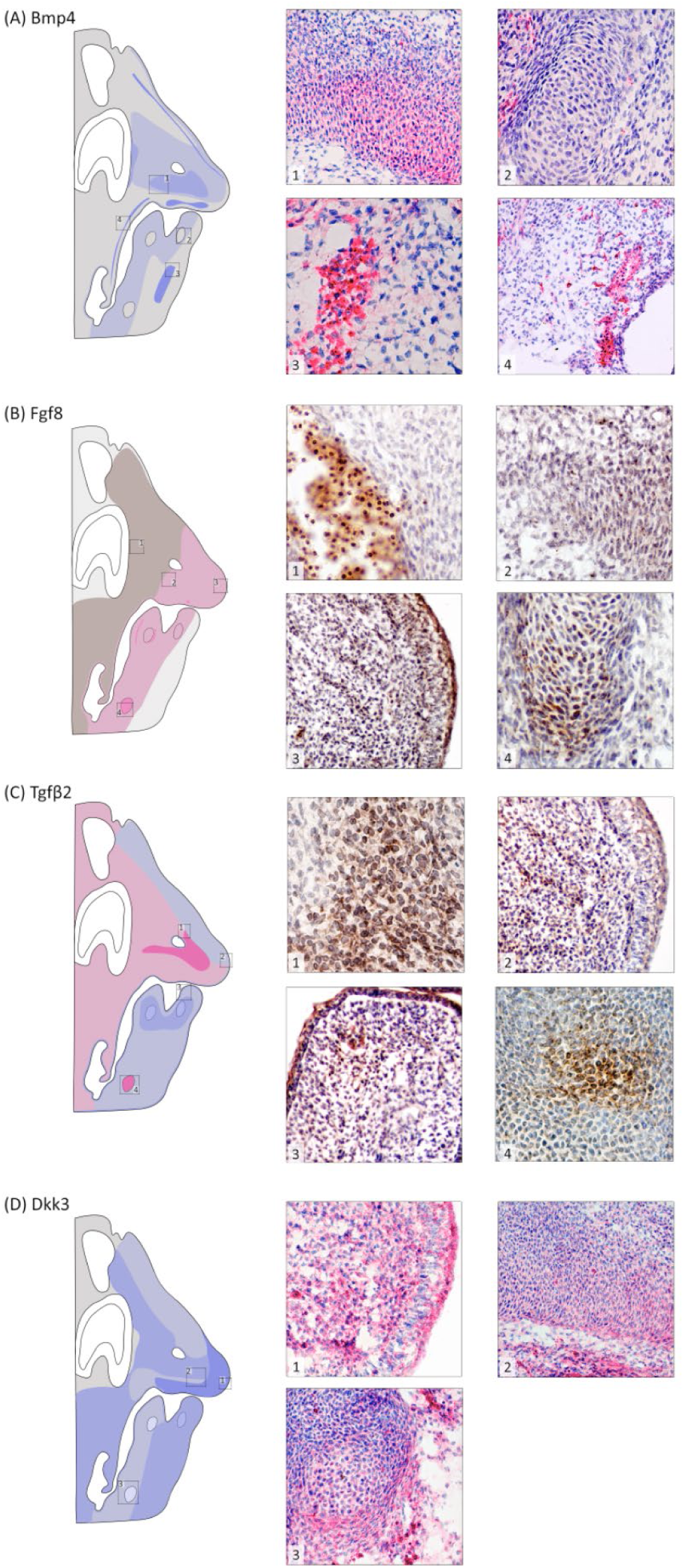
Bmp4, Fgf8, Tgfβ2, and Dkk3 expression at stage HH32. (A) **Bmp4** shows scattered cytoplasmic expression in upper and lower jaws with (1) moderately strong expression in the large condensation of the maxillary prominence. (2) Emerging cartilage is free of expression. (3) Strong expression in a broad stripe of premyogenic cells in the mandibular prominence. (4) The oral epithelium shows weak expression throughout with stripes of strong expression along the oral cavity along the length of the palatine process associated with developing vasculature. (B) **Fgf8** expression becomes more widespread at this stage with overall weak cytoplasmic and perinuclear expression throughout. (1) Strong nuclear expression in hematopoietic cells with weak cytoplasmic expression in adjacent mesenchyme. (2) Region below nasal cavity of maxillary prominence shows weak cytoplasmic expression with pockets of perinuclear expression. (3) Tip of maxillary prominence shows weak to moderate expression in surface epithelium and the underlying mesenchyme with pockets of perinuclear expression throughout. (4) Strong and largely perinuclear expression in basihyal cartilage. (C) **TGF-β** shows generally weak cytoplasmic expression in the upper and lower beak prominences with areas of strong perinuclear expression. (1) Condensed cells surrounding the nasal cavity area show strong perinuclear expression with posterior cells showing weak cytoplasmic and perinuclear expression while anterior cells show only weak cytoplasmic expression. (2) Weak cytoplasmic expression continues to tip of maxillary prominence. (3) Weak expression in the mesenchyme at the tip of the mandibular prominence with moderate epithelial expression. (4) Strong cytoplasmic and perinuclear expression in the basihyal cartilage formation. (D) **Dkk3** shows cytoplasmic expression overall that varies from weak to strong depending on location. (1) Strong expression at the tip of the maxillary prominence. (2) Bands of strong, weak and moderate expression (from bottom to top) associated with developing vascular regions, loosely organized mesenchymal cells, and the dominant central condensation of the maxillary prominence, respectively. (3) Strong expression at the edge of the basihyal cartilage with weak expression within. Very strong expression in hematopoietic cells (visible on rightmost side). See Figure 2 for key to anatomical features and expression patterns. Stain colors as in Figure 4.

**Figure 7.**
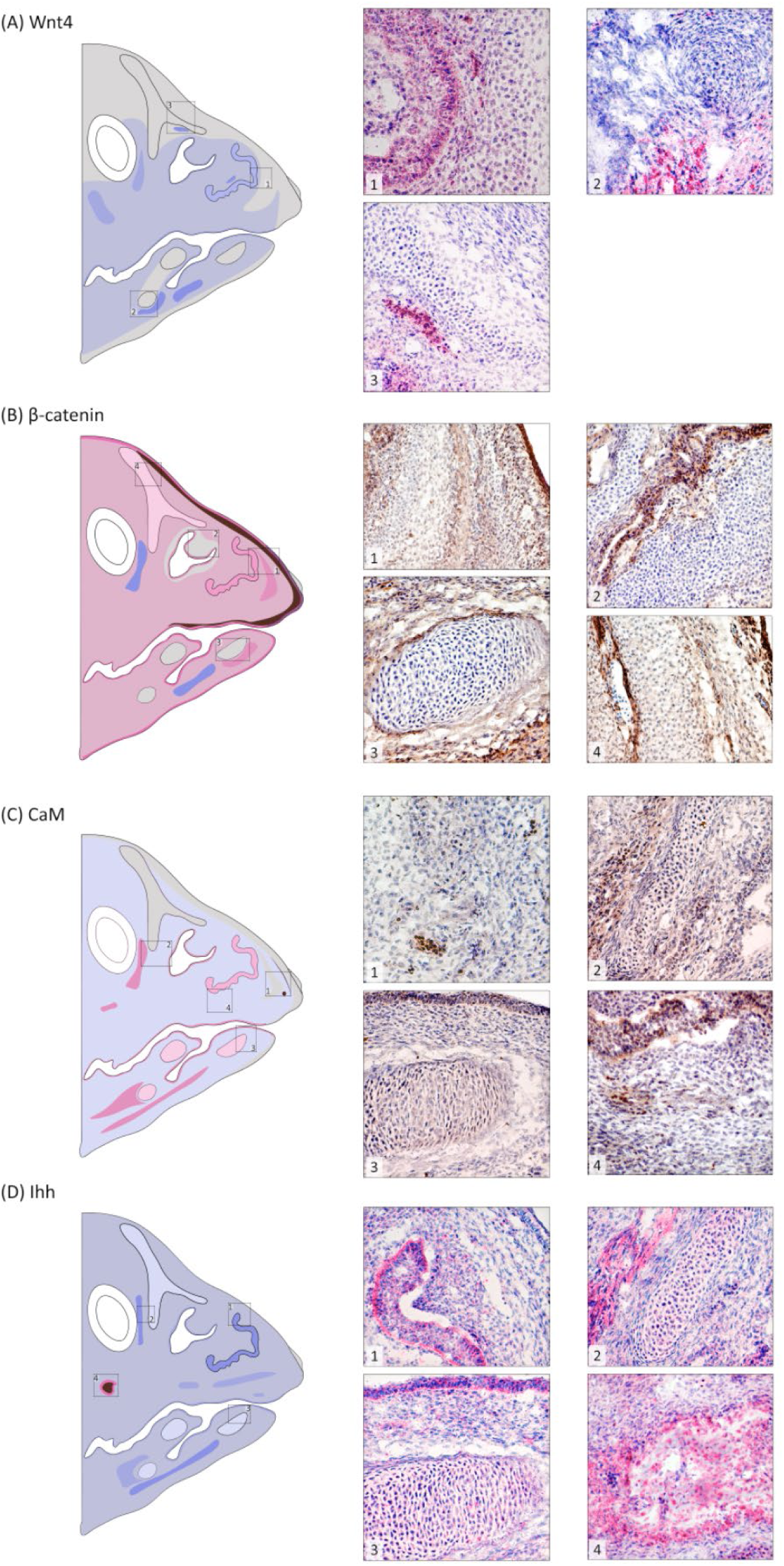
Wnt4, β-catenin, CaM, and Ihh expression at stage HH36. (A) **Wnt4** shows relatively weak cytoplasmic expression overall with tissue-specific areas of moderate to strong expression. (1) Moderate expression in the nasal epithelium with weak expression in surrounding cartilage. (2) No expression in paraglossal cartilage and strong expression in myogenic region directly below it. (3) Nasal cartilage free of expression, but a pocket of hematopoietic cells shows strong expression. (B) **β-catenin** shows overall weak cytoplasmic and nuclear expression in mesenchyme of both jaws with (1) strong membranal expression in the epithelium of the upper jaw tip. Underlying mesenchyme shows alternating bands of strong and weak nuclear/cytoplasmic expression. In the center of this region osteoprogenitor cells show moderate cytoplasmic and nuclear expression. (2) Expression is absent in the nasal cartilage but moderate in the nasal epithelium. (3) Meckel’s cartilage is free of expression with weak to moderate expression in surrounding mesenchyme. (4) Upper region of nasal cartilage exhibits some weak cytoplasmic and nuclear expression. There is continued absence of expression in hematopoietic cells. (C) **CaM** shows weak cytoplasmic expression in mesenchyme of both jaws with (1) continued absence of expression at the beak tip and strong nuclear expression in hematopoietic cells. (2) Weak, mostly cytoplasmic, expression in the anterior portion of the upper jaw with moderate cytoplasmic and nuclear expression in emerging muscle. (3) Weak perinuclear and cytoplasmic expression in the Meckel’s cartilage and its surrounding mesenchyme. Moderate cytoplasmic and perinuclear expression in the overlying oral epithelium. (4) Moderate nuclear and cytoplasmic expression in the nasal epithelium. Emerging osteogenic region (center) shows weak to moderate expression in a few cells. (D) **Ihh** shows overall weak cytoplasmic expression in the mesenchyme of both jaws. (1) Strong cytoplasmic expression in the nasal epithelium. (2) Weak cytoplasmic expression in the nasal cartilage with strong expression in adjacent muscle. (3) Weak expression in the Meckel’s cartilage with moderate expression in the overlying oral epithelium. (4) There is strong cytoplasmic and perinuclear expression in the emerging osteogenic region toward the back of the upper jaw. See Figure 2 for key to anatomical features and expression patterns. Stain colors as in Figure 3.

**Figure 8.**
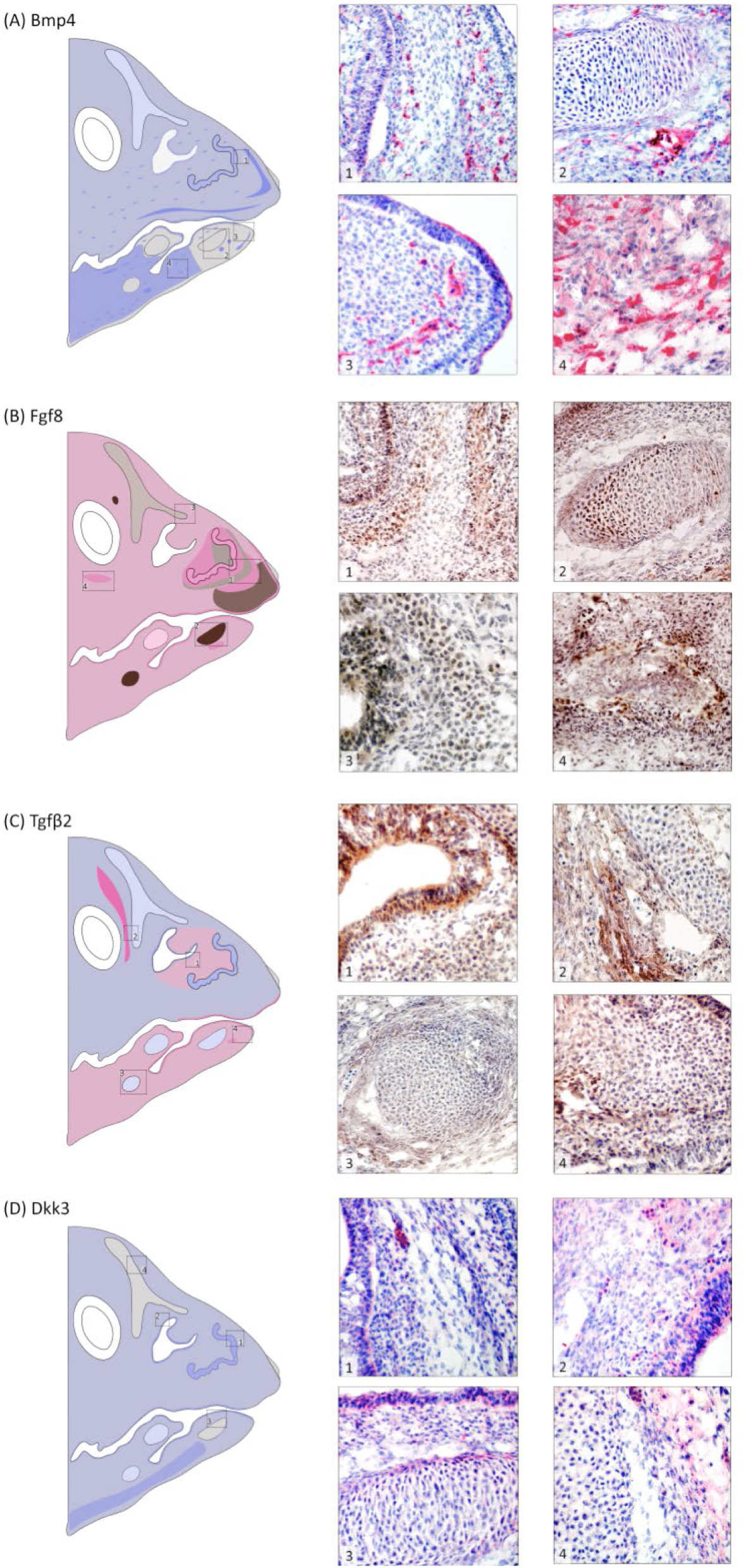
Bmp4, Fgf8, Tgfβ2, and Dkk3 expression at stage HH36. (A) **Bmp4** shows scattered cytoplasmic expression in upper and lower jaws. (1) Weak expression in the nasal epithelium and typical scattered expression in the adjacent mesenchyme with strong spots of expression interspersed with cells completely free of expression. (2) Meckel’s cartilage is free of expression with strong expression in perivascular and hematopoietic cells. (3) Tip of mandibular prominence largely free of expression with scattered regions of strong expression. (4) Posterior region of the lower jaw shows moderate to strong expression throughout. (B) **Fgf8** expression shows overall weak cytoplasmic and nuclear expression throughout with areas of stronger expression in emerging tissue. (1) Strong nuclear and cytoplasmic expression in the nasal epithelium with bands of nuclear and cytoplasmic expression in adjacent mesenchyme. (2) Meckel’s cartilage shows strong nuclear expression, particularly at the periphery. (3) Moderate nuclear to perinuclear expression in the nasal cartilage. (4) Strong nuclear and cytoplasmic expression at an emerging osteogenic region in posterior maxillary prominence. (C) **TGF-β** mesenchymal expression in upper and lower jaws differ in subcellular localization with upper beak showing largely cytoplasmic expression while lower beak showed predominantly cytoplasmic and perinuclear expression. (1) Nasal epithelium shows moderate, largely cytoplasmic expression while adjacent cartilage shows more perinuclear expression. (2) Weak cytoplasmic expression in upper nasal cartilage with a band of strong perinuclear and cytoplasmic expression in muscle. Hematopoietic cells show no expression. (3) Weak cytoplasmic and perinuclear expression in the basihyal cartilage with moderately strong perichondrogenic expression. (4) Weak expression in epithelium of the mandibular prominence with regions of mesenchyme showing moderate cytoplasmic and perinuclear expression. (D) **Dkk3** shows weak cytoplasmic expression throughout the upper and lower beak mesenchyme. (1,2) Moderate expression in the nasal epithelium compared to surrounding mesenchyme. (3) Meckel’s cartilage shows no expression with weak expression in adjacent mesenchyme. Overlying oral epithelium shows moderate expression. (4) No expression in upper nasal cartilage, weak expression in surrounding mesenchyme and strong expression in hematopoietic cells (visible in upper center). See Figure 2 for key to anatomical features and expression patterns. Stain colors as in Figure 4.

**Figure 9.**
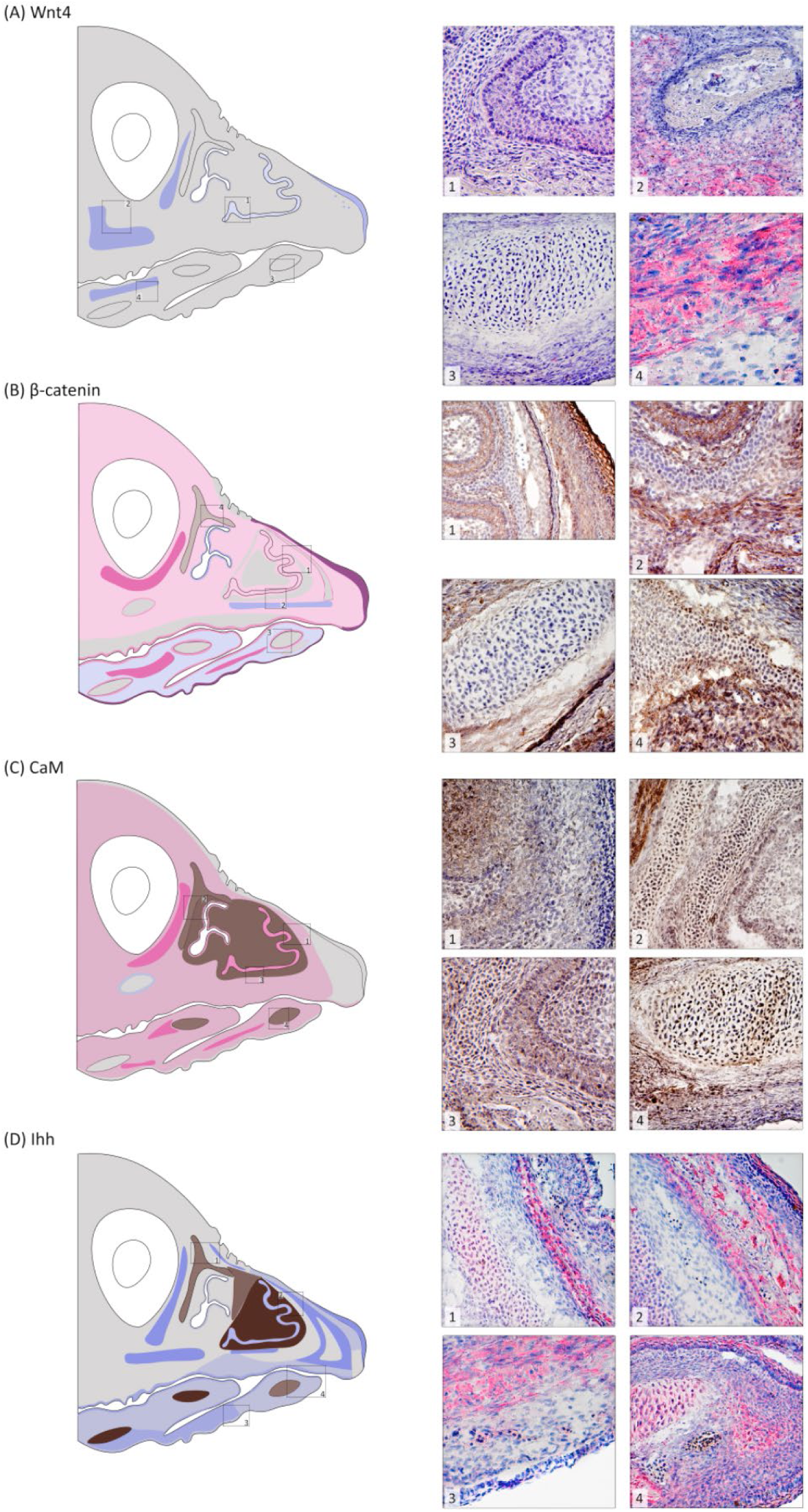
Wnt4, β-catenin, CaM, and Ihh expression at stage HH39. (A) **Wnt4** shows limited cytoplasmic expression in upper and lower beak mesenchyme. (1) Weak expression in the nasal epithelium with no expression in underlying osteogenic region. (2) Both osteogenic center and periosteogenic region are free of expression. Strong expression in developing muscle below this area. (3) No expression in Meckel’s cartilage and surrounding mesenchyme. (4) Emerging muscle in lower jaw shows strong expression. (B) **β-catenin** shows overall weak cytoplasmic and nuclear expression in mesenchyme of the upper beak and mostly cytoplasmic expression in lower beak. (1) Bands of weak to moderate cytoplasmic and nuclear expression are associated with emergence of distinct tissue types in upper jaw. Cartilage shows mostly nuclear expression, while the nasal epithelium shows mostly cytoplasmic and the mesenchyme shows both. Central areas of the emerging osteogenic region are largely free of expression while the edges show a strong band of expression. Surface epithelium shows strong membranal expression. (2) Similar tissue-specific patterns as in 1 with additional views of moderate expression in emerging muscle and no expression in hematopoietic cells. (3) Meckel’s cartilage remains free of expression with weak to moderate expression in surrounding mesenchyme. A thin band of cells surrounding the osteogenic envelope shows strong nuclear and cytoplasmic expression. (4) Upper region of nasal cartilage shows moderate cytoplasmic and nuclear expression. (C) **CaM** shows weak cytoplasmic and nuclear expression in mesenchyme of upper and lower jaw with exception of the upper beak tip. (1) Nuclear and cytoplasmic expression in nasal epithelium with largely nuclear expression in surrounding cartilage. No expression in surface epithelium and its underlying mesenchyme. (2) Strong nuclear expression in nasal cartilage and strong cytoplasmic and nuclear expression in muscle. (3) Similar patterns of tissue expression as in 1 with moderate nuclear and cytoplasmic expression in osteogenic region (bottom third). (4) Strong nuclear expression in Meckel’s cartilage with moderate cytoplasmic and nuclear expression in surrounding mesenchyme. Surface epithelium is free of expression. (D) **Ihh** shows overall limited expression in upper beak mesenchyme with more extensive expression in the lower beak mesenchyme. (1) Weak cytoplasmic expression in the nasal cartilage surrounded by mesenchyme with no expression. An emerging osteogenic region shows strong cytoplasmic expression. (2) Nasal cartilage with weak perinuclear expression. Adjacent mesenchymal and hematopoietic cells are free of expression while the osteogenic region shows strong cytoplasmic expression in both the center and periphery. (3) Moderate cytoplasmic expression in muscle of the lower beak. (4) Moderate perinuclear expression in the Meckel’s cartilage, with moderate expression in the overlying oral epithelium. Mesenchymal expression is mostly weak with pockets of stronger expression in osteogenic regions. See Figure 2 for key to anatomical features and expression patterns. Stain colors as in Figure 3.

### Protein coexpression within stages

#### Stage HH29/30

At this stage, β-catenin showed nuclear expression in both the upper and lower beak, particularly in the epithelium of the anterior edge of the maxillary and mandibular prominences (Figure 3). Fgf8 showed the most limited expression (Figure 4), while CaM and Fgf8 displayed nuclear expression in condensations (Figures 3C-2, 4B-2), and Ihh showed strong cytoplasmic expression in these areas (Table 1). Tgfβ2, β-catenin and Dkk3 expressed strongly at the periphery of condensations compared to the other proteins (e.g. Figures 3B-, 4C-3, 4D-1). Wnt4, Ihh, and Fgf8 had higher expression in the upper beak, whereas other proteins were more evenly expressed across the upper and lower beaks (Figures 3, 4). Bmp4 was most strongly expressed in linear tracts in the subepithelial mesenchyme along the ventral margin of both the maxillary and mandibular prominences (e.g. Figure 4A-2,3). Broader tracts of expression in other proteins (e.g. β-catenin, CaM, and Dkk3) in the lower jaw aligned with pre-myogenic regions (Figures 3, 4).

**Table 1.**
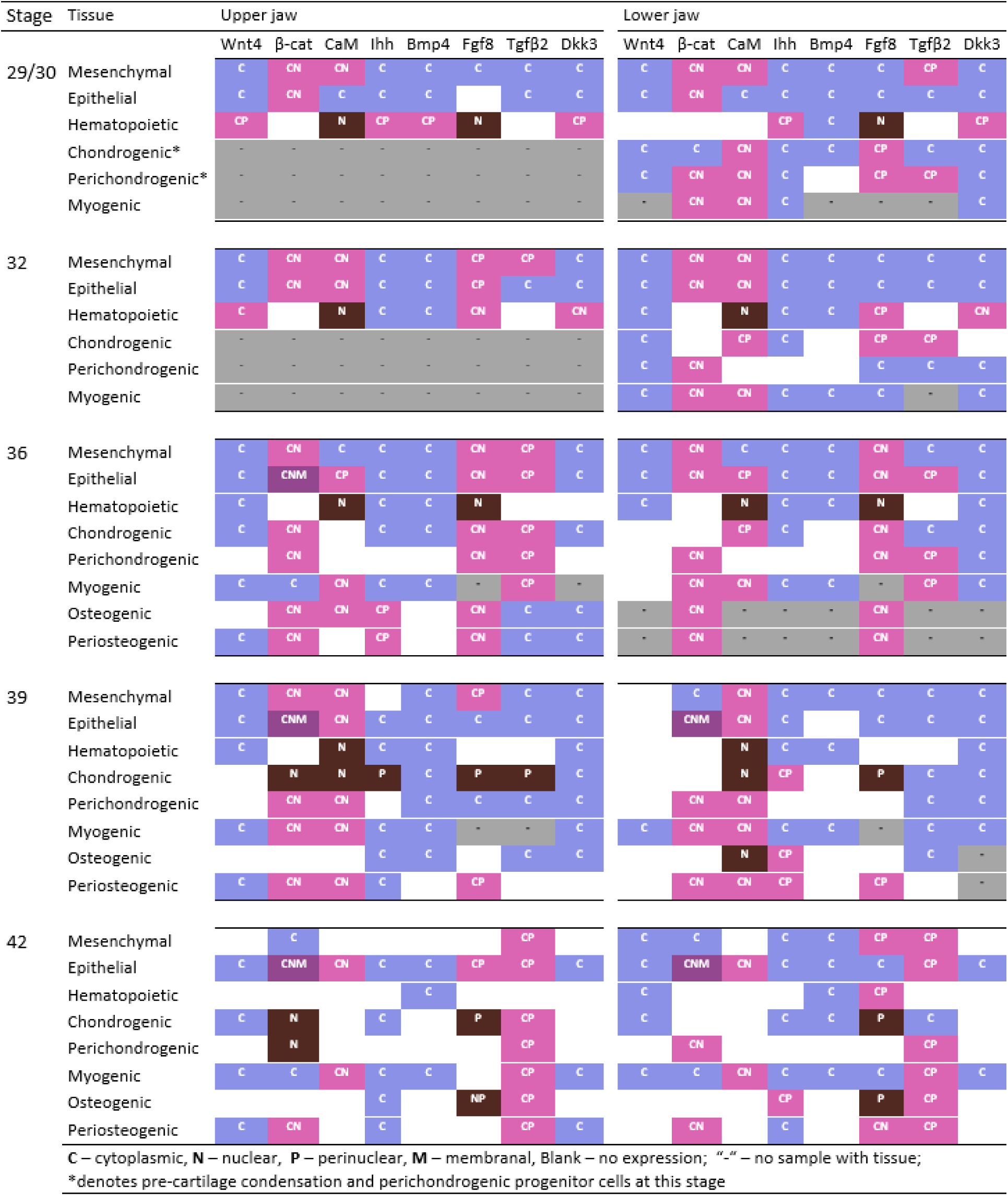
Cellular localization of protein expression across developmental stages in upper and lower jaw.

#### Stage HH32

Several proteins showed distinct expression patterns around the nasal cavity region of the upper jaw (Figures 5, 6). For example, Wnt4, Ihh, Bmp4, Tgfβ2 all had elevated expression in the region directly below the nasal cavity, particularly in an area of highly condensed cells (Figures 5A-1, 5D-2, 6A-1, 6C-1). Notably, Tgfβ2 showed strong perinuclear expression here. In addition to the greater expression in the centrally located condensation, several proteins also showed differential expression along the anterior-posterior axis of the upper beak. For example, Dkk3 and β-catenin were progressively higher toward the upper beak tip (Figures 5B-1, 6D-1), whereas Bmp4, Ihh, CaM and Wnt4 showed reduced expression at the beak tip (Figures 5, 6). In the lower jaw, Wnt4, Ihh, Bmp4, and Fgf8 were absent toward the ventral margin and in the surface epithelium (e.g. Figure 5A-3, 5D-3); only β-catenin and Dkk3 maintained strong expression in this region (Figures 5B-3, 6D-3).

**Figure 10.**
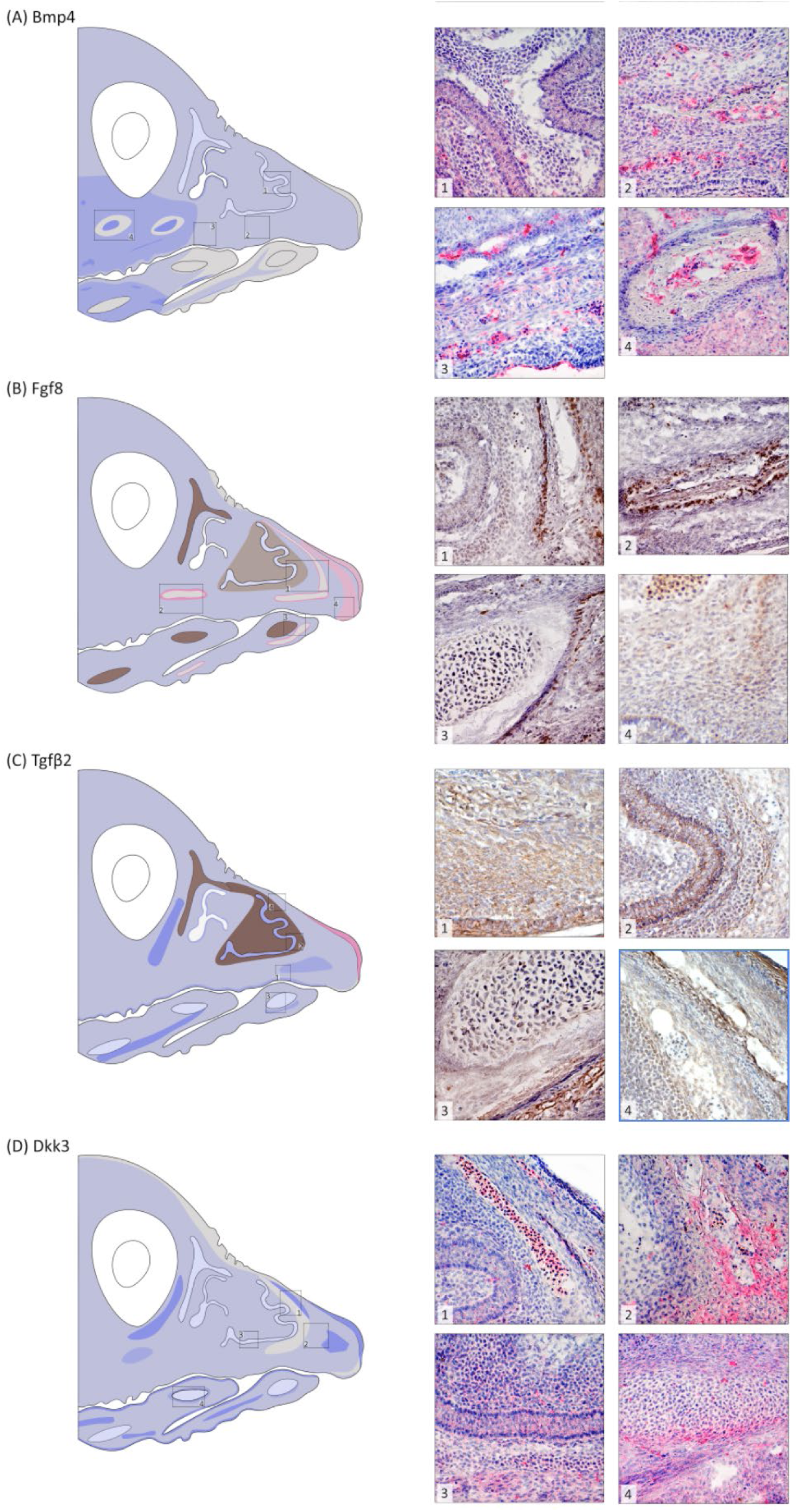
Bmp4, Fgf8, Tgfβ2, and Dkk3 expression at stage HH39. (A) **Bmp4** shows weak cytoplasmic expression in the upper jaw with a gradient of no expression, weak expression and moderate expression from the tip to the back of the lower jaw. (1) Weak expression in the nasal epithelium with some scattered expression in the adjacent cartilage. (2) Weak expression in upper jaw mesenchyme with strong expression in emerging osteogenic region. (3) Weak expression in muscle of lower jaw with areas of strong expression directly adjacent and at the bottom edge of the tongue. Epithelial tissue is largely free of expression. (4) An osteogenic region of the upper beak shows strong expression in the center with a conspicuous ring of no expression directly surrounding it. Away from this osteogenic region, mesenchyme shows weak cytoplasmic expression. (B) **Fgf8** expression shows weak cytoplasmic expression in most of the upper and lower jaw mesenchyme. (1) Weak cytoplasmic expression in the nasal epithelium with bands of weak perinuclear expression in adjacent cartilage. (2) Strong perinuclear and cytoplasmic expression in the periphery of an osteogenic region of the upper beak. (3) Moderate perinuclear expression in the Meckel’s cartilage with very weak cytoplasmic expression in the surrounding mesenchyme with heightened expression in an emerging osteogenic region. (4) The upper beak tip is the only mesenchymal region to show strong perinuclear and cytoplasmic expression. (C) **TGF-β** mesenchymal expression in upper and lower beaks shows largely cytoplasmic expression overall. (1) Strong cytoplasmic expression in muscle with cytoplasmic and perinuclear expression in cartilage. Hematopoietic cells are free of expression. (2) Strong cytoplasmic expression in nasal epithelium with strong perinuclear expression in the adjacent cartilage. (3) Meckel’s cartilage shows very weak cytoplasmic expression with moderately strong expression in adjacent osteogenic region (bottom right). (4) Strong perinuclear expression in nasal cartilage and in nearby osteogenic region with only weak cytoplasmic expression in mesenchyme and surface epithelium. (D) **Dkk3** shows weak cytoplasmic expression throughout the upper and lower beak mesenchyme. (1) Weak expression in the nasal epithelium, no expression in nasal cartilage and moderate expression in hematopoietic cells. (2) Strong expression in osteogenic region of the emerging premaxillary bone. (3) Weak to moderate expression in nasal epithelium and its surrounding cartilage in more posterior regions of the upper beak. (4) Paraglossal shows some weak expression with stronger expression in the perichondrogenic region. See Figure 2 for key to anatomical features and expression patterns. Stain colors as in Figure 4.

**Figure 11.**
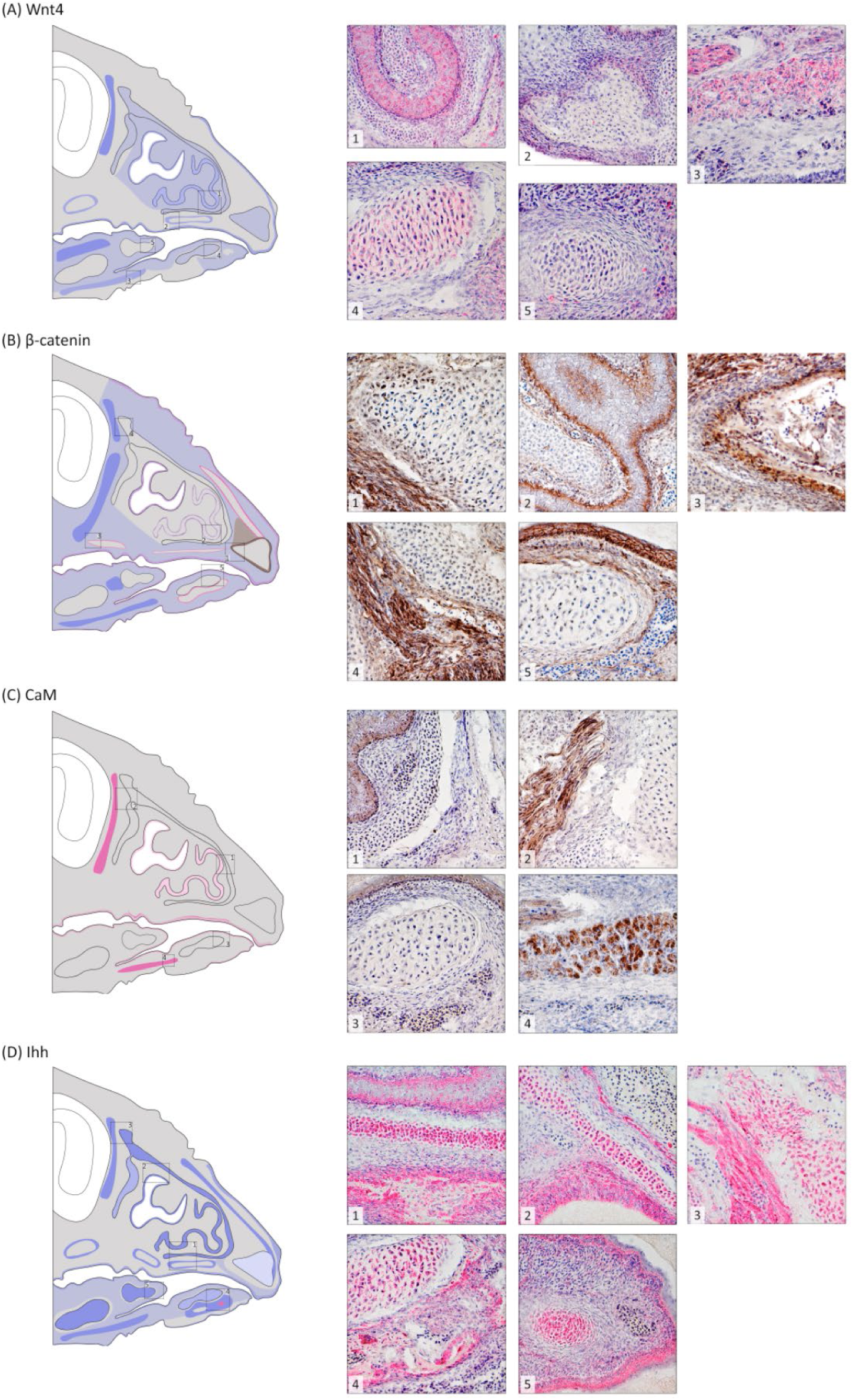
Wnt4, β-catenin, CaM, and Ihh expression at stage HH42. (A) **Wnt4** is largely absent in upper beak mesenchyme with limited expression in the lower beak mesenchyme. (1) Moderate expression in the nasal epithelium and weak expression in surrounding cartilage. (2) Osteogenic center is free of expression while periosteogenic area shows weak expression. (3) Moderate expression in muscle of lower jaw while surrounding mesenchyme is largely free of expression. (4) Weak cytoplasmic expression in the Meckel’s cartilage. (5) Paraglossal cartilage is largely free of expression. (B) **β-catenin** shows overall weak cytoplasmic expression in mesenchyme of the upper and lower beak. (1) Premaxillary cartilage center is free of expression with strong nuclear expression at edges. (2) Nasal epithelium shows moderate expression at its base, while surrounding cartilage is free of expression. (3) Muscle (top left) and osteogenic regions show strong cytoplasmic and nuclear expression. (4) Upper nasal cartilage is largely free of expression with some weak expression at edges. (5) Meckel’s cartilage is free of expression and strong cytoplasmic and membranal expression continues in epithelial tissues. Hematopoietic cells show no expression. (C) **CaM** shows no expression in mesenchyme of upper and lower jaw. (1) Weak nuclear and cytoplasmic expression in nasal epithelium with no expression in surrounding cartilage or osteogenic region. (2) Strong nuclear and cytoplasmic expression in muscle. (3) No expression in Meckel’s cartilage with weak cytoplasmic and nuclear expression in overlying epithelium. (4) Strong cytoplasmic and nuclear expression in muscle of lower beak. (D) **Ihh** shows overall limited expression in upper beak mesenchyme with more extensive expression in the lower beak mesenchyme. (1) Strong cytoplasmic expression in the nasal epithelium, nasal cartilage and in an underlying periosteogenic region. (2) Nasal cartilage and epithelium show strong expression while hematopoietic cells show no expression. (3) Strong expression in muscle and nearby upper nasal cartilage. (4) Strong cytoplasmic expression in the Meckel’s cartilage, with moderate expression in surrounding mesenchyme. Osteogenic region shows both cytoplasmic and perinuclear expression at its center. (5) Strong expression in paraglossal cartilage with a notable absence of expression in perichondrogenic area. Moderate expression in epithelium of tongue. See Figure 2 for key to anatomical features and expression patterns. Stain colors as in Figure 3.

**Figure 12.**
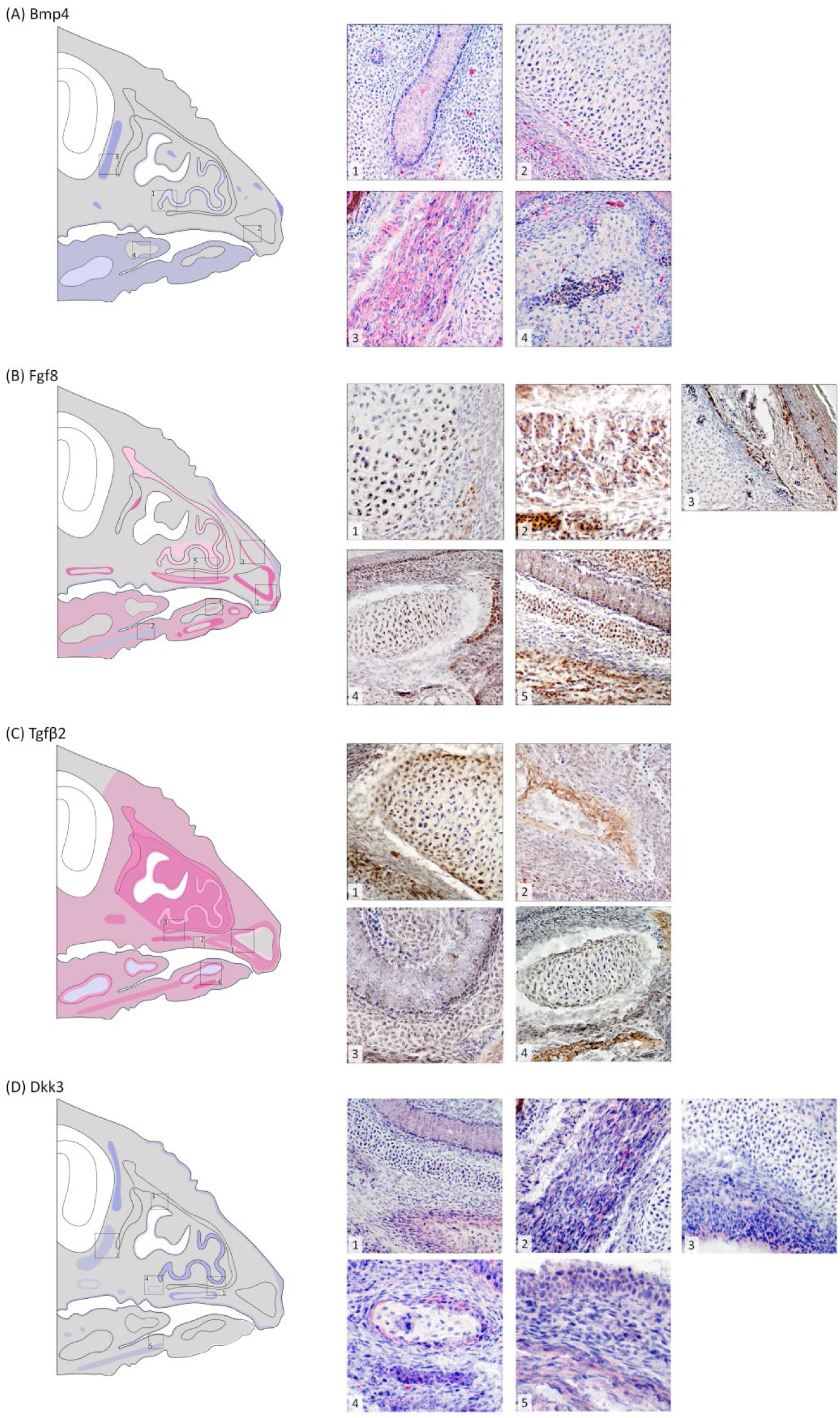
Bmp4, Fgf8, Tgfβ2, and Dkk3 expression at stage HH42. (A) **Bmp4** shows no mesenchymal expression in the upper beak with weak to moderate expression in the lower beak. (1) Weak expression in the nasal epithelium. (2) No expression in premaxillary cartilage and weak expression in the underlying epithelium. (3) Strong expression in upper jaw muscle. Adjacent cartilage is largely free of expression. (4) Osteogenic region of the Meckel’s shows no expression around edges or the center with limited scattered expression in the mesenchyme and the oral epithelium. (B) **Fgf8** expression shows overall no expression in most of the upper jaw mesenchyme with weak perinuclear and cytoplasmic expression in the lower jaw. (1) Strong perinuclear expression at the edges of the premaxillary cartilage and perinuclear/cytoplasmic expression at the beak tip. (2) Moderate cytoplasmic expression in muscle of the lower beak with strong expression in hematopoietic cells. (3) Strong periosteogenic expression in the upper beak, weak expression in the surface epithelium, and no expression in anterior nasal cartilage. (4) Meckel’s cartilage shows strong nuclear expression in center with no expression in perichondrogenic area. (5) Weak cytoplasmic and nuclear expression in the nasal epithelium, weak nuclear expression in the nasal cartilage and strong nuclear expression in adjacent osteogenic area. (C) **Tgfβ2** mesenchymal expression in upper and lower beaks shows weak cytoplasmic and perinuclear expression overall. (1) Strong expression in perichondrogenic region of pre-maxillary with additional weak expression at cartilage edges. (2) Weak to moderate expression in mesenchyme and osteogenic region of upper beak. (3) Expression varies from weak in epithelium, to moderate in surrounding cartilage, to strong in muscle. (4) Meckel’s cartilage is largely free of expression with some weak cytoplasmic, especially around edges. (D) **Dkk3** shows limited expression throughout the upper and lower beak mesenchyme. (1) Weak expression in the nasal epithelium, no expression in nasal cartilage and weak expression in the periosteogenic region. (2) Weak expression in muscle. (3) Weak to no expression at edge of nasal epithelium. (4) Weak to moderate periosteogenic expression in upper beak. (5) Weak expression in oral epithelium and muscle of lower jaw. See Figure 2 for key to anatomical features and expression patterns. Stain colors as in Figure 4.

#### Stage HH36

Protein expression was most spatially distinct at this stage, with some proteins (e.g., β-catenin, Fgf8) retaining strong expression throughout the upper and lower beak, while others (e.g., Wnt4 and CaM) were weakly expressed toward the beak tip and in the surface epithelium of the upper beak (Figures 7, 8). Most proteins showed strong expression in differentiating tissues such as myogenic areas and nasal epithelium but had only weak expression in mesenchyme (Figures 7, 8). Several proteins diverged in expression pattern across the oral, nasal and surface epithelium. For example, CaM and Wnt4 were absent in the surface epithelium but highly expressed in the oral and nasal epithelia (Figure 7A-1, 7C-3,4). Other proteins, such as Ihh, Tgfβ2 and Bmp4, were weakly expressed in surface epithelia (e.g. Figures 7D-1, 8A-1), but strongly expressed in the oral epithelium (e.g. Figures 7D-3, 8C-4). β-catenin and Bmp4 were largely absent from chondrogenic regions (Figures 7A-3,4, 8A-2) but exhibited weak expression in the upper nasal cartilage (e.g. Figure 7B-4). In contrast, CaM, Ihh, Fgf8, and Tgfβ2 were expressed in chondrogenic areas, all showing nuclear or perinuclear expression (Figures 7C-3, 7D-2,3, 8B-2,3, 8C-1,3).

#### Stage HH39

β-catenin, Tgfβ2, and CaM maintained strong expression in the mesenchyme while the other proteins show more limited mesenchymal expression (Figures 9, 10). Strong expression of β-catenin at the beak tip shifted from nuclear to membranal (Figure 9B-1). Wnt4 was absent in chondrogenic regions (Figure 9A-3) while β-catenin and Bmp4 were expressed only in the upper portion of the nasal cartilage (e.g. Figure 9B-4). Dkk3 expression was relatively weak in the mesenchyme but strong in osteogenic, myogenic and perichondrogenic regions (Figure 10D). In contrast, strong expression within chondrogenic areas persisted for CaM and Ihh (Figure 9C-2,3,4, 9D-1,2,4). Tgfβ2 showed strong perinuclear expression in the upper beak nasal cartilage (Figure 10C-1), but mostly weak cytoplasmic expression in the lower beak (Figure 10C-3). Wnt4 was expressed strongly in the mesenchyme surrounding a posterior osteogenic area but showed no periosteogenic or central osteogenic expression (Figure 9A-2). β-catenin and Fgf8 displayed strong periosteogenic expression (Figures 9B-3, 10B-1,2) and Fgf8 also showed weak scattered expression within these regions especially in the premaxillary osteogenic region (Figure 10B-4). Ihh was strongly expressed in the periosteogenic and central osteogenic regions (Figure 9D-1,2). CaM and Tgfβ2 show weak to moderate expression in and around osteogenic regions (Figures 9C-3,4, 10C-4). Both Dkk3 and Bmp4 were strongly expressed in osteogenic centers with Bmp4 showing a striking lack of expression in the immediate periosteogenic areas (Figure 10A-4, 10D-2)). Notable differences in expression patterns across the upper and lower jaw emerge for β-catenin which began to show less nuclear expression in the mesenchyme of the lower jaw, whereas expression of CaM, Fgf8 and Bmp4 formed pronounced gradients along the anterior-posterior axis of the upper beak (Figures 9B, 9C, 10A, 10B).

#### Stage HH42

Only β-catenin and Tgfβ2 had widespread mesenchymal expression in the upper beak at this stage with other proteins confined to tissue-specific expression. However, in the mesenchyme of the lower beak Wnt4, Fgf8, Bmp4, and Ihh maintained strong expression (Figures 11, 12). Wnt4 was weakly expressed in premaxillary, Meckel’s and nasal cartilage, but was absent from paraglossal and basihyal cartilages (e.g. Figure 11A-1,4,5). CaM and Dkk3 were absent in cartilage (Figures 11C-1,2,3, 12D-1,2,3) while Fgf8 showed limited perinuclear expression in the premaxillary and Meckel’s cartilage (Figure 12B-1,4). Tgfβ2 and β-catenin were expressed at the edge of the premaxillary cartilage and in the nasal septum (Figures 11B-1,4, 12C-1,3). Bmp4 had only scattered expression at the edge of the premaxilla and basihyal cartilage (e.g. Figure 12A-2). Ihh was strongly expressed in chondrogenic regions of the lower beak and the nasal cartilage (Figure 11D). Most proteins at this stage showed moderate to strong expression in myogenic areas (Figures 11, 12). Wnt4 and Dkk3 were expressed in the mesenchyme surrounding osteogenic areas (Figures 11A-2, 12D-4). β-catenin and Fgf8 continue to show strong periosteogenic expression (Figures 11B-3, 12B-3,4) and Fgf8’s expression within these areas was largely confined to posterior regions of the upper jaw and most of the lower jaw (Figure 12). Ihh was strongly expressed in the periosteogenic and central osteogenic regions (Figure 11D-1,4). CaM was absent from osteogenic regions (Figure 11C-13) while Tgfβ2 was strongly expressed in and around these areas (Figure 12C-2,4). Bmp4 showed more moderate and scattered expression in osteogenic centers (Figure 12A-4).

## Discussion

By mapping core regulatory proteins across five stages that span chondro- and osteogenesis, our atlas (Figures 3-12) provides the first integrative view of histological and subcellular contexts of protein expression during late ontogeny of the passerine beak. Many of the regulatory proteins studied here are known to interact in complex and context-dependent ways^59–61^. By assessing their co-expression patterns, we can identify interactions among pathways that may be inhibitory or synergistic. We can also identify spatial and temporal contexts of expression that may be particularly crucial for overall patterning and tissue differentiation.

### Shared and divergent expression patterns

The observed patterns of protein expression are largely consistent with known molecular interactions among craniofacial gene pathways (Figure 1). For example, β-catenin showed strong nuclear expression at the beak tip throughout stages 29-36 consistent with its role in maintaining a proliferative growth zone during premaxillary outgrowth^58^. As growth slowed in later stages, β-catenin shifted to strong membranal expression in the surface epithelium consistent with its function in adherens junctions and regulation of epithelial structure^61^. This shift alters the dynamics of underlying mesenchymal cells, for example by promoting denser packing and reduced motility (cell jamming), thereby creating conditions that could favor mechanical activation of Tgfβ2 in the extracellular matrix. It could also modulate the availability of the other extracellular-matrix associated ligands studied here, including Bmp4, Fgf8, Ihh, and Wnt4^62,63^. Dkk3 also shows strong expression at the beak tip at stage HH32, which corroborates its role in regulating premaxillary bone growth downstream of β-catenin and Tgfβ2 signaling^58,64^. Prior work inferring these interactions also showed upregulation of β-catenin and Tgfβ2 signaling at the beak tip^58,64^ which differs from our findings as we did not find strong expression of the Tgfβ2 ligand at the beak tip. However, this prior work focused on *TGFβIIr* mRNA expression, not the ligand itself. Instead, we found strong perinuclear expression of Tgfβ2 in the condensation of the upper beak suggesting that ligand production is upregulated more centrally, possibly leading to increased extracellular matrix storage, allowing these cells to function as a source of latent TGFβ2 that can act paracrinely on receptor-rich cells at the beak tip.

Additional pathway interactions suggested by prior work (Figure 1) were evident in the early formation of chondrogenic areas. For example, expression of Ihh within pre-chondrogenic condensations and Bmp4 and Tgfβ2 in surrounding regions was consistent with feedback loops in which Ihh/Gli and Bmp/Tgfβ–Smad pathways (Figure 1) jointly regulate chondrocyte proliferation and hypertrophy in cartilage growth plates^65,66^. Likewise, strong nuclear β-catenin in perichondrogenic mesenchyme (e.g. Figure 2B-1) is consistent with its role in preventing premature or excessive chondrocyte differentiation and setting up boundaries between cartilage core areas and surrounding tissues^67^.

We found moderately strong nuclear expression of CaM in precartilage condensations. While other studies have found Ca²⁺/CaM-dependent regulation of nuclear targets in cartilage^10,68,69^, direct visualization of nuclear CaM in early craniofacial condensations has, to our knowledge, not been documented. However, its expression in conjunction with strong Ihh, weak β-catenin and Tgfβ2 expression within pre-cartilage condensations is consistent with a cell population that is transitioning from pre-condensation mesenchyme toward early chondrogenesis. The overlap of high Dkk3 and β-catenin in the perichondrogenic region has not been reported before and may indicate a role for Dkk3 in boundary formation, perhaps as a mechanism of buffering against excessive Wnt signaling. Dkk3 is often an antagonist of Wnt/β-catenin signaling^47^ however, its role is context-dependent and in this context, it may work with β-catenin to fine-tune Wnt activity. Further work is needed to elucidate Dkk3’s role as a potential modulator of Wnt–Bmp/Tgfβ balance at cartilage and bone boundaries. Together, these patterns support many of the canonical points of intersection highlighted by previous studies, while revealing potentially novel contributions of CaM and Dkk3 in coordinating formation of early condensations.

We observed strong expression of Ihh, Fgf8, and Tgfβ2 in chondro- and osteogenic tissues, reflecting their known roles in coordinating cartilage differentiation and bone matrix formation^27,34,70,71^. Ihh expression was strong in both central and periosteogenic regions of ossifying tissue, consistent with its dual function in promoting chondrocyte hypertrophy^72,73^ and mediating signaling between cartilage and adjacent osteogenic fronts^74^. Perichondrogenic and periosteogenic regions were also enriched in Tgfβ2 and Fgf8, consistent with their roles in the regulation of differentiation at developing cartilage and bone fronts. Bmp4 was notably absent from periosteogenic margins with strong expression in osteogenic centers at stage HH39 (Figure 9). This pattern is consistent with Bmp4’s role in promoting osteogenic differentiation and matrix deposition and suggests that its expression may be inhibited at the edges of osteogenic regions to delineate ossification boundaries. In contrast, β-catenin was strongly expressed in periosteogenic regions and at the margins of the premaxillary cartilage at stage HH42. This latter pattern may reflect its role in promoting differentiation of mesenchymal progenitor cells into osteoblasts while blocking their progression toward chondrocytes^75,76^. Wnt4 was more peripherally distributed around osteogenic centers, suggesting a modulatory or signaling feedback role in refining the boundaries of bone-forming regions. Together, these patterns suggest that osteogenesis in the zebra finch beak arises from a coordinated interface between signaling-active boundary cells and differentiating cores, emphasizing that cross-pathway feedback is essential in establishing bone and cartilage differentiation while maintaining boundaries between distinct tissue compartments.

At the subcellular level IHH and TGFβ2 showed consistent perinuclear and FGF8 nuclear expression within chondro- and osteogenic regions, indicating active signaling within differentiating cells, while Bmp4 remained cytoplasmic and was instead associated with osteogenic patterning. Fgf8 has not previously been known to act in the nucleus during craniofacial development and is largely viewed as a signaling molecule^77^. However, recent studies have proposed intranuclear functions of Fgf proteins, including Fgf8^78,79^ and have found that some Fgfs have nuclear localization signals allowing them to migrate into the nucleus and interact with other transcription factors to modulate the expression of target genes^80,81^. These studies, together with our findings, suggest an expanded view of Fgf8’s role during craniofacial morphogenesis as a potential transcription coregulator.

#### Spatial and temporal contexts of expression

The dynamics of regulatory protein expression (Figures 3-12) indicate that stage HH32 may be a crucial inflection point for tissue compartmentalization. With the exception of CaM and β-catenin, most proteins at this stage showed elevated expression around the central condensation of the upper beak (Figures 5 and 6). This spatially localized expression suggests the nasal cavity may be acting as a focal signaling node as upper-beak compartments become established. Moreover, Dkk3, Fgf8 and β-catenin showed elevated expression toward the beak tip suggesting that proximal-distal patterning is becoming established and that tissues are beginning to adopt more position-specific regional identities. These developmental events directly follow the fusion of the craniofacial prominences which occurs at stage HH28-29 in the chick^82,83^. Together, these findings suggest that stage HH32 may be a key transition point where regional protein expression begins to define the morphogenetic compartments of beak structure.

Developmental dynamics of the lower beak is understudied^84^ with available work pointing to pronounced heterochrony related to distinct developmental origins of upper and lower beak tissues, as well as compensatory interactions between them^21^. We found that progression of tissue differentiation and patterning of the upper and lower beak differed strongly across developmental stages corroborating prior work^85,86^. By stage HH29/30 pre-cartilage condensations had formed in the lower beak with strong expression of several proteins in pre-myogenic and pre-cartilage condensations (Table 1), whereas the upper beak exhibited a delayed and more centralized pattern of activity emerging at stage HH32. After stage HH32, patterns of expression across the upper and lower beak became more similar with subcellular localization of protein expression broadly similar in chondro- and osteogenic tissue of both beaks.

A notable pattern at stage HH36 was divergence in expression of the oral, surface and nasal epithelia (e.g. see Figure 5C and D). The oral epithelium often showed strong and coordinated patterns of expression in the upper and lower beak that were distinct from the surface epithelium suggesting it may be a coordinating center for growth as well as a source of positional information for tissue differentiation. Sánchez Moreno and Badyaev^87^ showed that, in the house finch (*Haemorhous mexicanus*), prior to tissue transformations, early arriving neural crest-derived mesenchymal (NCM) cells transiently match protein profiles of the overlying epithelium producing distinct boundaries that anchor mesenchymal cell condensations. Our findings similarly suggest that epithelial tissue may be a primary source of morphogenetic signals that establish patterning in the avian beak.

## Conclusions

In this study, we bridge molecular and histological perspectives, providing a foundation for understanding how signaling dynamics are translated into morphological form. By documenting cellular colocalization of protein expression in its tissue-level context we reveal points of pathway convergence for proliferative, chondrogenic, and osteogenic phases of avian beak development. By mapping eight core regulatory proteins across five developmental stages and multiple tissue types, our study extends previous work that examined these factors in isolation or at early stages, offering the first systematic depiction of how signaling activity changes across the maturation of the beak. Bridging molecular and histological perspectives provides a synthesis for the timing and specification of tissue transformations, not just in later stages of beak development but also across the upper and lower beak, two current gaps in the field. Ultimately, our map will provide a foundation for comparative studies of craniofacial development as well as evolution of the avian beak.

### Experimental procedures

#### Embryo Collection, Staging and Histology

Zebra finches were housed in a large semi-outdoor (roofed with open wire mesh on two sides) walk-in aviary (4.72L x 1.52W x 2.44H m) at the University of Arizona in accordance with the Institutional Animal Care and Use Committee’s guidance and approval (IACUC: 16-111). In this colony, which houses from 80-100 zebra finches at any given time, individuals are allowed to pair freely and breeding is closely monitored (see^88^ for details). Throughout breeding birds receive finch mix seed and water *ad libitum*, in addition to grit, calcium supplements, prophylactic coccidiosis medication (preventative maintenance dosage, semi-weekly), weekly millet spray, and twice weekly dietary supplements of spinach and hard-boiled egg. Nest boxes in the colony were checked daily for fresh eggs, which were marked during laying to keep track of laying order and developmental timing. Collected embryos were viewed under a dissecting scope to confirm embryo stage^56^. Five stages spaced throughout the final third of development were chosen for sectioning (HH29/30, HH32, HH36, HH39, and HH42; *n* = 10). Many of the embryos at stage 29/30 were collected midway between these two stages and so are lumped for analysis. After staging, embryos were fixed and stabilized using PaxGene (Pre-Analytix, Switzerland) and dehydrated in 30% sucrose in PBS. The heads of embryos were flash frozen in dimethylbutane and placed in OCT media (Fisher Scientific, Itasca, IL) before cryosectioning at 10-12 µm. Slides were stored at -80°C until immunohistochemistry (IHC). Before IHC we identified the approximate midline of each embryo to standardize location for tissue staining. Eight sections per stage were stained with hematoxylin and eosin using standard procedures to differentiate among tissue types and visualize tissue transformation across developmental time^63^ (H&E; U. Rochester MC).

#### Immunohistochemistry

IHC was optimized for eight antibodies (Table S1) using positive and negative control tissue (esophagus, kidney, esophagus, testes, and small intestine) collected and sectioned as described above (Table S1, Figure S1) as well as isotype controls (Cell Signaling Technologies, Danvers, MA; see^62,87^ for details).

For immunostaining, we used anti-β-catenin (610153, 1:16,000, BD Transduction Laboratories), anti-CaM (sc-137079, 1:15, Santa Cruz Biotechnology), anti-Wnt4 (ab91226, 1:800; Abcam), anti-Tgfβ2 (ab36495, 1:800, Abcam), anti-Bmp4 (ab118867, 1:100, Abcam), anti-Ihh (ab184624, 1:100, Abcam), anti-Dkk3 (ab214360, 1:100, Abcam), and anti-Fgf8 (89550, 1:50, Abcam) antibodies using methods described previously^89^. Validations confirming specificity of stains are in Badyaev^62^. Reactions were visualized with either diaminobenzidine (DAB, Elite ABC HRP Kit, PK-6100, Vector Labs) or Vector Red Alkaline Phosphatase substrate and Vectastain ABC-AP Kit (AK-5000, Vector Labs). Slides were counterstained with Mayer’s hematoxylin, blued in ammonium water, dehydrated, cleared, and mounted with Permount mounting medium (Fisher Scientific) prior to analysis. Three slides, each containing four tissue sections (12 sections per embryo) were run with the following grouped antibodies: i) β-catenin, Fgf8, Tgfβ2 and no primary control, ii) Bmp4, Wnt4, Ihh and no primary control, and iii) Dkk3 and no primary control, and CaM and no primary control. The control sections on each slide did not receive the primary antibody during IHC but were otherwise treated the same as other sections.

#### Photomicroscopy and Image Assessment

IHC staining was visualized with brightfield optics (Olympus microscope, Japan). Pictures were taken with a microscope digital camera (Olympus DP74) and the Olympus cellSens software. All images were taken at standardized settings to ensure consistency across slides. Images were taken at low magnification (4x and 10x) to summarize patterns throughout the upper and lower beak and at high magnifications (20x and 40x) to show cellular level details. For each stage, two individuals were sectioned and stained, and within each individual staining patterns were assessed for consistency across multiple IHC runs (2-4 runs per individual). We then created a consensus staining pattern from these images, documenting presence/absence of expression, cellular localization, and types of tissues stained. Qualitative assessments of high, medium and low levels of expression were based on comparison of relative expression levels within sections only. A consensus map of expression was determined based on the maximum area stained across different individuals and sections and the most inclusive subcellular localization designation was used for Table 1.

## Acknowledgements

We thank Fatima Bravo for help with IHC assays and Neha Varshney for assistance with histological identification. The Badyaev and Duckworth lab groups provided helpful feedback that improved this manuscript. This project was supported by grants from the National Science Foundation (IBN-0218313 and DEB-1754465) to A.V.B.

## Data availability statement

upon acceptance, all data will be made available in a publicly accessible repository.

## Ethics approval statement

All animal procedures were conducted in accordance with the Institutional Animal Care and Use Committee (IACUC) of the University of Arizona (# 16-111)

---

**Figure S1.**
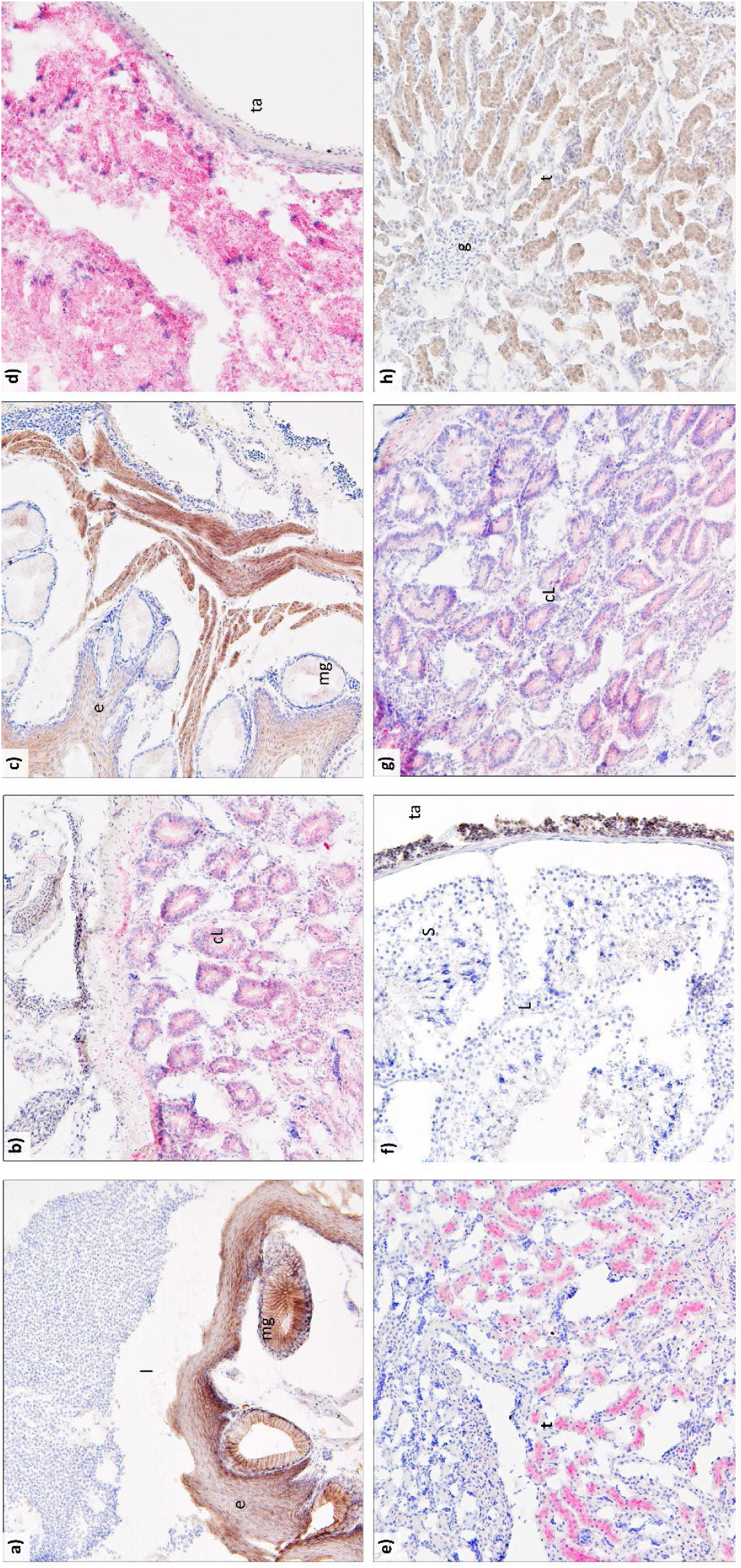
IHC Optimization in positive and negative control tissue. (a) β-catenin expression (brown stain) in esophagus: positive expression in epithelium and mucous glands, negative expression in the lumen. (b) BMP expression (red stain) in kidney: positive expression in tubules, negative expression in glomeruli. (c) CaM expression (brown stain) in esophagus: positive expression in epithelium, negative expression mucous glands and lumen. (d) Dkk3 expression (red stain) in testes: positive expression in Sertoli cells and Leydig cells, negative expression in tunica albuginea. (e) Ihh expression (red stain) in small intestine: positive expression in crypts of Lieberkuhn, negative expression in the lamina propria. (f) Fgf8 expression (brown stain) in testes: positive expression in tunica albuginea, negative expression in Sertoli cells and Leydig cells. (g) Wnt4 expression (red stain) in small intestine: positive expression in crypts of Lieberkuhn, negative expression in the lamina propria. (h) TGFβII expression (brown stain) in kidney. Positive expression in tubules, negative expression in glomeruli. cl = crypt of Lieberkuhn, e = epithelium, g = glomerulus, mg = mucous gland, L = Leydig cells, l = lumen, S = Sertoli cells, t= tubule, ta = tunica albuginea, lm = lamina propria

**Table S1.** Antibodies used for IHC.

| **Protein of interest** | **Antibody** | **Manufacturer** | **Dilution** |
| --- | --- | --- | --- |
| $\beta$ -catenin | Anti-β-catenin<br>(from mouse) | BD Transduction Labs<br>(#610153, San Jose, CA) | 1:16,000 |
| Bone morphogenic<br>protein 4 (Bmp4) | Anti-Bmp4<br>(from mouse) | Abcam Biotechnology<br>(#118867, Cambridge,<br>MA) | 1:100 |
| Calmodulin (CaM) | Anti-CaM<br>(from mouse) | Santa Cruz<br>Biotechnology (#137079,<br>Dallas, TX) | 1:20 |
| Dickkopf-3 (Dkk3) | Anti-Dkk3<br>(from rabbit) | Abcam Biotechnology<br>(#214360, Cambridge,<br>MA) | 1:90 |
| Fibroblast growth<br>factor 8 (Fgf8) | Anti-Fgf8<br>(from mouse) | Abcam Biotechnology<br>(#89550, Cambridge,<br>MA) | 1:50 |
| Indian hedgehog<br>(Ihh) | Anti-Ihh (from<br>rabbit) | Abcam Biotechnology<br>(#184624, Cambridge,<br>MA) | 1:100 |
| Transforming<br>growth factor beta<br>type 2 (Tgfβ2) | Anti-TGFβII<br>(from mouse) | Abcam Biotechnology<br>(#36495, Cambridge,<br>MA) | 1:800 |
| Wingless type 4<br>(Wnt4) | Anti-Wnt4<br>(from rabbit) | Abcam Biotechnology<br>(#91226, Cambridge,<br>MA) | 1:800 |

